# Site-specific arrangement and structure determination of minor groove binding molecules in self-assembled three-dimensional DNA crystals

**DOI:** 10.1101/2023.10.10.561756

**Authors:** Chad R. Simmons, Alex Buchberger, Skylar J.W. Henry, Alexandra Novacek, Nour Eddine Fahmi, Tara MacCulloch, Nicholas Stephanopoulos, Hao Yan

## Abstract

The structural analysis of guest molecules in rationally designed and self-assembling DNA crystals has proven elusive since its conception. Oligonucleotide frameworks provide an especially attractive route towards studying DNA-binding molecules by using three-dimensional lattices with defined sequence and structure. In this work, we site-specifically position a suite of minor groove binding molecules, and solve their structures via x-ray crystallography, as a proof-of-principle towards scaffolding larger guest species. Two crystal motifs were used to precisely immobilize the molecules DAPI, Hoechst, and netropsin at defined positions in the lattice, allowing us to control occupancy within the crystal. We also solved the structure of a three-ring imidazole-pyrrole-pyrrole polyamide molecule, which sequence-specifically packs in an anti-parallel dimeric arrangement within the minor groove. Finally, we engineered a crystal designed to position both netropsin and the polyamide at two distinct locations within the same lattice. Our work elucidates the design principles for the spatial arrangement of functional guests within lattices and opens new potential opportunities for the use of DNA crystals to display and structurally characterize small molecules, peptides, and ultimately proteins of unknown structure.

## INTRODUCTION

The foundational goal of structural DNA nanotechnology has been to rationally design three-dimensional (3D) lattices made entirely of DNA, and use them to precisely orient unknown guest species in order to solve their atomic structure.^1^ Since this proposal, significant advances in structural biology have made high resolution structure determination of macromolecules more readily achievable. Nevertheless, DNA crystals remain attractive scaffolds for the arrangement of molecules in 3D space with atomic precision once the structure of the DNA lattice itself is known. Much like other porous materials—such as metal organic frameworks (MOFs), covalent organic frameworks (COFs), or porous coordination polymers, which have served as effective scaffolds for small organic molecules, nanoparticles, metal ions, and biomolecules^2-6^—DNA crystals have seen a range of applications in recent years. These include crystal frameworks for catalysis, as storage or separation devices, or for delivering cargo using non-canonical interactions.^7-16^

Self-assembling DNA crystals are constructed by using standard Watson-Crick base pairing rules along with multi-arm branched motifs (immobile Holliday junctions)^17-20^ and sticky ends that facilitate the assembly of the oligonucleotides in 3D space. These frameworks can be used to precisely orient molecules or oligonucleotide binding proteins requiring specific binding sequences, or small DNA and RNA aptamers, with the potential to solve their structure using x-ray crystallography. To this end, an expanding toolbox of these lattices containing a wide variety of crystal symmetries, tunable pore volumes and sizes, and programmable sequences have been developed, including a recently reported 3D double crossover motif that demonstrated that the crystals were viable for binding of Hoechst 33342 for structural studies.^9,15,21-30^

The synthesis of small molecules that can mimic the sequence-specific binding of certain proteins to DNA has been an attractive pursuit for decades, especially to design artificial transcription factors for therapeutics.^31-36^ Dervan and colleagues proposed that polyamide molecules could preferentially bind to DNA sequences within the minor groove (MG) of DNA, and differentiate between G•C/C•G and A•T/T•A pairs.^37-40^ This was first accomplished with the crystal structure of netropsin, which is a polyamide molecule known to have antibiotic and antiviral activity,^41^ containing two aromatic *N*-methylpyrrole (Py) rings, bound to an AATT sequence in the MG of the duplex.^42,43^ Netropsin belongs to a larger family of small molecules that bind to A/T rich regions of the MG, including cell staining dyes such as 4’,6-diamidino2-phenylindole (DAPI) and Hoechst 33342 and 33258.^44^

Based upon netropsin, it was proposed that molecules replacing one of the Py rings with a N-methylimidazole (Im) moiety could potentially recognize a G•C substitution.^42,45^ A three-ring polyamide ImPyPy-Dp was designed to explicitly recognize a TGTCA binding sequence, by stacking in an anti-parallel manner within the MG. Specifically, when paired with the imidazole on one end, its counterpart pyrrole carboxamide stacks on the opposing molecule, thereby allowing specified recognition of a G•C pairing, whereas when reversed, a C•G pairing is targeted.^39,46,47^ The anti-parallel (ImPyPy)_2_-DNA packing with 2:1 stoichiometry within the MG was initially unexpected until the NMR structure of distamycin (PyPyPy) was solved,^48^ followed by the NMR structure of the (ImPyPy)_2_-DNA;^39,46,47^ however, to our knowledge, the crystal structure of (ImPyPy)_2_ bound to TGTCA has never been solved.

Herein, we describe the use of programmable, self-assembled DNA crystal lattices as host systems for minor groove binders (MGBs) DAPI, Hoechst 33342, and netropsin (Figure 1). We introduced a consensus binding sequence (AATT) into previously designed crystals, obtained co-crystals of these lattices with the bound MGBs, and solved their structures using x-ray crystallography for direct comparison to existing structures (Figure 1a-c). In addition, we show that the molecules can be strategically positioned at defined MGs, while controlling the occupancy of the guest within the crystal, with prescribed spatial arrangement of the molecules in the lattice, dependent upon crystal symmetry. Lastly, we programmed a unique (ImPyPy)_2_-DNA binding sequence (TGTCA) into the DNA lattice (Figure 1d). The structure of the molecule was solved at both a single MG position, and then in the presence of netropsin at the alternate MG, to high resolution (2.45 Å), confirming the molecular interactions and sequence selectivity of the original polyamide design.

**Figure 1.**
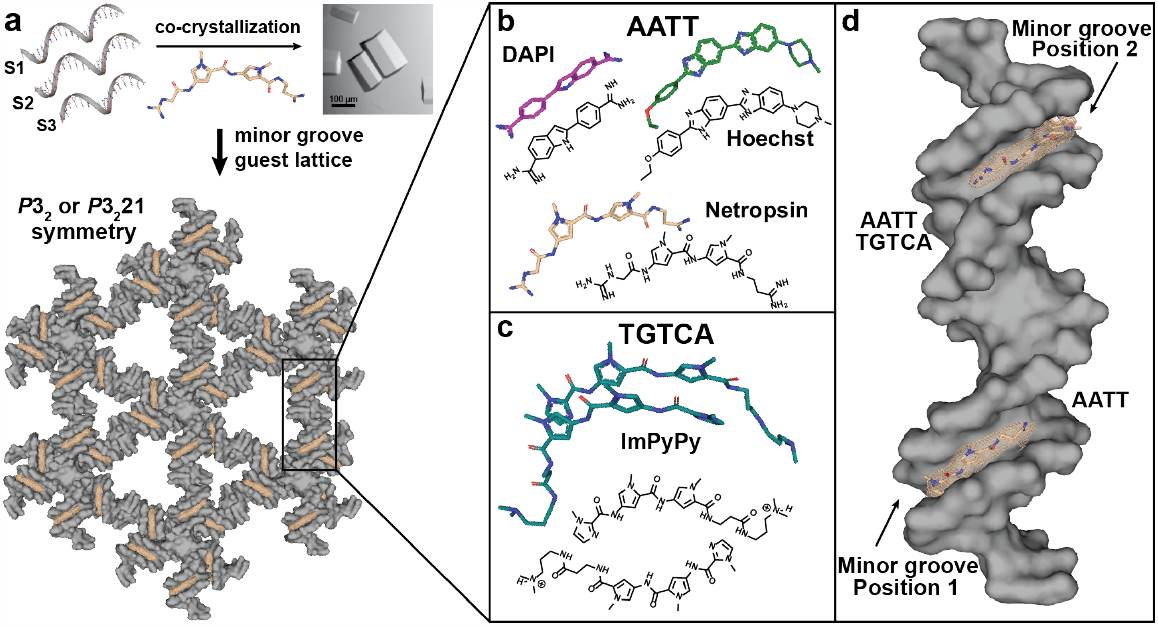
Schematic of the assembly of the DNA lattices co-crystallized in the presence of minor groove binding molecules. (a) Three component oligonucleotides (S1-3) co-crystallized in the presence of minor groove binding molecules (netropsin is shown here) are used to assemble three-dimensional lattices for two crystal motifs (4x5 and 4x6). The fundamental component to the lattice is a 21-bp duplex (rectangular box) with complementary 2-bp sticky ends and two Holliday junction crossover points that allow the lattice to form into continuous 3D layered helical arrays that can hold a binding sequence specific for the molecule at each minor groove position in the crystal. The “empty” electron density (tan) results from the presence of the molecule bound to the DNA prior to model building; (b) Co-crystals were prepared with three unique AATT binding molecules (DAPI (purple), Hoechst (green), and netropsin (tan)), and their structures are described in detail in the work. The chemical structures corresponding to each respective molecule are also shown; (c) An anti-parallel three-ring imidazole-pyrrole-pyrrole (ImPyPy-β-Dp) polyamide molecule (teal) which binds sequence specifically to a TGTCA sequence; and (d) Each component duplex (gray surface) comprising the lattice in (a) contains two minor grooves that can bear an AATT sequence specific to the molecule at either Position 1 on the 5’ end of S2, Position 2 on the 3’ end, or at both available minor grooves. The 2Fo – Fc electron density contoured at s = 1.5 (tan) for the bound netropsin molecule results when the AATT sequence is placed at both minor grooves in the co-crystals. All carbon molecules are colored according to the assigned color of each molecule, with oxygen (red) and nitrogen (blue) shown according to atom type.

## RESULTS AND DISCUSSION

### Experimental design

In this work, we employed two DNA crystal motifs (4x5 or 4x6) requiring three constituent oligonucleotides: (S1; chain A) containing 4 repeats of 5 or 6 bases which forms crossovers at each repeat tethering each duplex within the lattice; (S2; chain B) a linear 21-base strand; and (S3; chain C & D) which is a 16-base crossover strand containing complementary sequences to S1 and S2 on either end of each duplex, resulting in the assignment of two chains (Supplementary Figure 1). Each 21-bp duplex is flanked by complementary 2-base sticky ends which cohere to form continuous linear layers (Figure 1a), with each Holliday junction containing the “J1” sequence.^18,19^ The MGs were programmed with a favored AATT binding sequence and we selected the MGBs DAPI, Hoechst 33342, and netropsin for co-crystallization (Figure 1c) because they bind to AT-rich regions with reported Kd values of 1.8 nM, 38 nM, and 1.0 x nM, respectively.^49-51^ Crystals were acquired with sparse matrix screening, and subsequently optimized (Figure 1a & b, Supplementary Figures 2 & 3 and Tables 1-4).

Each DNA duplex within the scaffold provides two unique MGs where we could site- and sequence-specifically arrange each molecule at a single position (Pos1), at the alternate MG (Pos2)—resulting in crystals with 50% of all available MGs occupied—and at both positions (BP), yielding fully saturated crystals (Figures 1b & Supplementary Figure 1). Lastly, we synthesized a modified version of a previously described ImPyPy-β-Dp (IPP) MGB molecule^38,39^ with binding sequence: TGTCA (Figure 1d). This sequence was placed at Pos2 in the 4x5 motif, whereas a subsequent construct was used containing AATT at Pos1, and TGTCA at Pos2 (Figure 1b), and co-crystals were obtained with both netropsin and IPP bound to their respective sequences (Supplementary Figures 4 & 5 and Table 5). The structures were solved using x-ray diffraction, all data collection and refinement statistics were summarized (Supplementary Tables 6-9), and the coordinates and structure factors were deposited in the Protein Data Bank.

### Crystal structures of MGBs in self-assembling DNA lattices

All co-crystals diffracted from 2.45-3.1 Å, and yielded structures containing positive difference density corresponding to each respective MGB. The molecules were subsequently independently built, followed by the calculation of polder omit maps,^52^ post-refinement, to provide additional evidence that the models were correct, and the resulting 2*F*_*o*_ *– F*_*c*_ and polder maps were superimposed to demonstrate their agreement (Supplementary Figures 6-11). The DAPI molecules were roughly planar, running parallel to the MG axis (Figure 2a-c). The atom positions for both the Hoechst and netropsin structures were well resolved with the density clearly indicating the tilting of the aromatic groups and curvature of the molecules along the contour of the MG (Figure 2d-i & Supplementary Figures 6-11). We note that without these details, the accurate determination of the directionality of the molecules would have been problematic. Lastly, the electrostatic contacts between the molecules and the minor groove were mapped (Figure 3; Supplementary Figure 12), and the original structures for DAPI, Hoechst, and netropsin (1D30, 129D, and 6BNA, respectively)^43,53,54^ were aligned to compare (RMSD) the structures in parallel (Supplementary Table 10 & Supplementary Figure 13). Although most of the previously described structures of these MGBs have resulted with much higher resolution detail, this study serves to describe how these molecules could be structured within the context of a self-assembled DNA lattice, even at moderate resolution.

**Figure 2.**
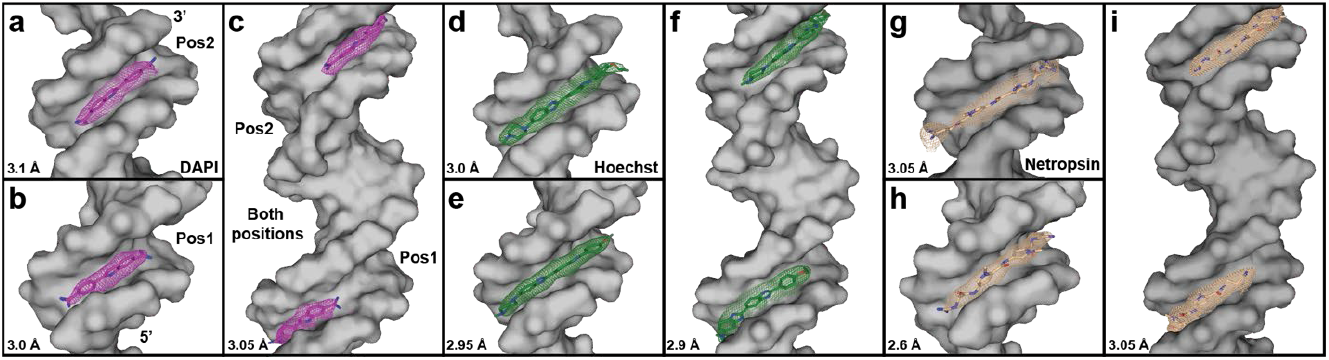
Programmable positioning of minor groove binders within the 4x5 DNA lattice. Structures for each individual minor groove position is represented for DAPI, Hoechst, and netropsin, and the 2Fo – Fc electron density for each MGB is shown. (a) Pos2 DAPI (purple) contoured at s = 1.2 (purple); (b) Pos1 DAPI contoured at s = 1.2; (c) Both positions DAPI contoured at s = 1.2; (d) Pos2 Hoechst (green) contoured at s = 1.0 (green); (e) Pos1 Hoechst contoured at s = 1.2; (f) Both positions Hoechst contoured at s = 1.0; (g) Pos2 netropsin (tan) contoured at s = 0.8 (tan); (h) Pos1 netropsin contoured at s = 1.1; and (i) Both positions netropsin with contoured at s = 1.6. The surface of the DNA (gray) for each respective site(s) is shown. The 5’ and 3’ ends of S2 (chain A) are labeled, and the resulting resolution of each respective crystal structure is indicated at the lower left of each panel.

**Figure 3.**
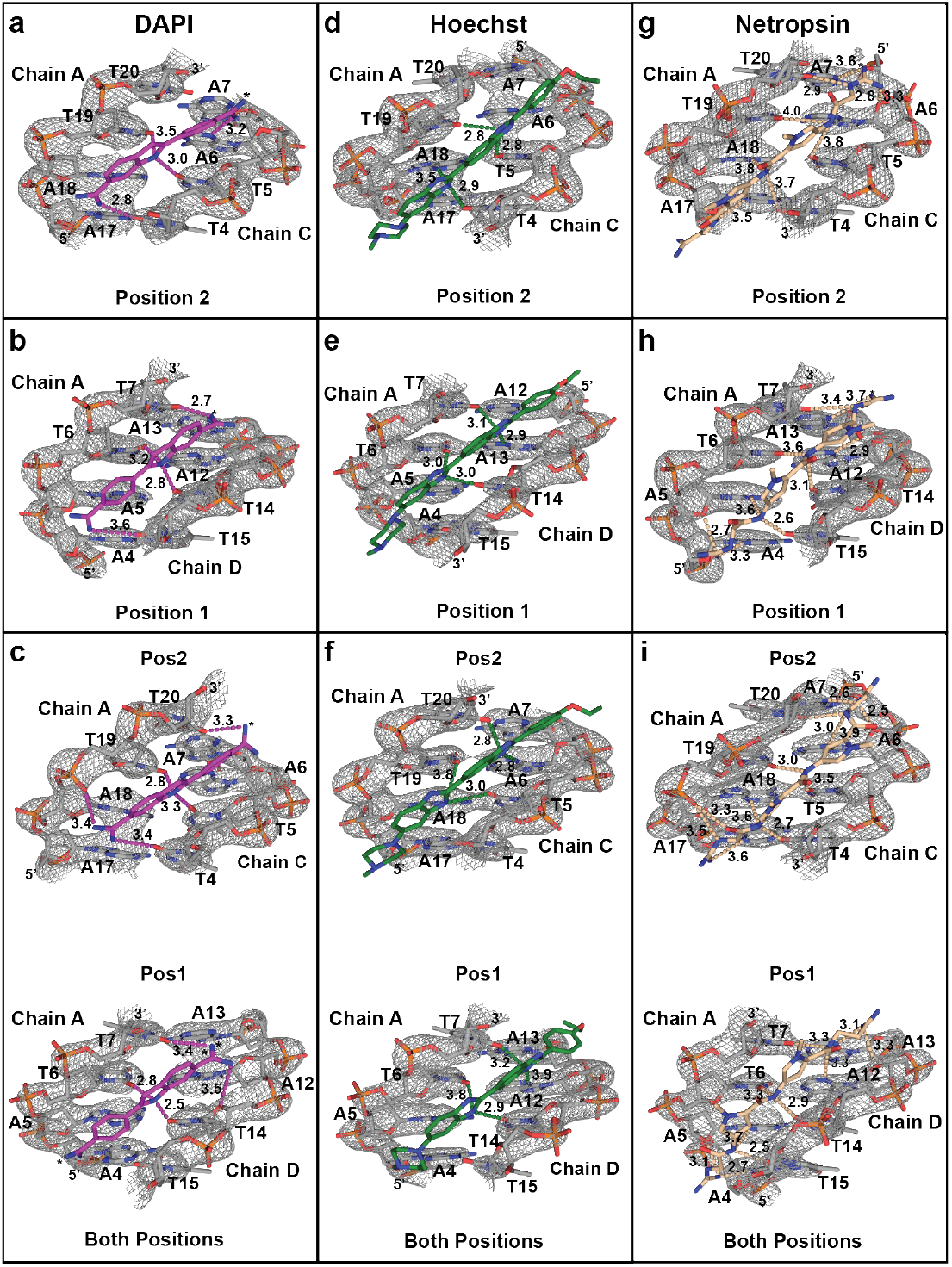
Structure and electrostatic coordination of DAPI, Hoechst, and netropsin to the AATT minor groove in the 4x5 lattices. The AATT sites of each minor groove (MG) within 2*F*_*o*_ *– F*_*c*_ electron density demonstrating the agreement of the map and model. The phosphate backbone, base stacking and pairing interactions, and purine-pyrimidine identities are well resolved. The chains and nucleotides are labeled with the 5’ and 3’ ends indicated to orient the direction of the sequence. All electrostatic contacts (dashes) and distances between the molecule and the DNA are shown, with asterisks indicating additional contacts outside the pocket. Distances of strong (2.5 Å) to weak (4.0 Å) interactions (dashes) were used. (a) Polar contacts of DAPI Pos2 MG contoured at s = 2.3., with additional contacts between chain A G8(O4’) and N3 of DAPI (3.6 Å); (b) DAPI Pos1 MG (σ = 2.6), with additional contacts between chain A G8(O4’) and N3 of DAPI (3.4 Å); (c) DAPI at both positions (BP) σ = 2.3 (Pos1) and 2.6 (Pos2), with additional contacts between chain D C14(O4’) and N4 of DAPI (3.5 Å), chain A A21(OP1) to DAPI-N2 (3.2 Å) and to A21(O4’) (3.0 Å), and DAPI-N2 and G8(O4’) of chain C (3.2 Å); (d) Hoechst 33342 at Pos2 at σ = 2.3; (e) Hoechst 33342 at Pos1 at σ = 2.4; (f) Hoechst 33342 at BP with DNA at Pos2 at σ = 2.8 and Pos1 at σ = 2.4; (g) netropsin Pos2 MG at σ = 2.5, with additional contacts between chain A A21(N3) and netropsin-N2 (2.8 Å); (h) netropsin Pos1 MG (σ = 2.5), with additional contacts between chain A C9(O4’) and netropsin-N10 (3.5 Å); (i) netropsin BP with Pos2 (σ = 3.1) and Pos1 (σ = 2.9), with additional contacts between chain A netropsin-N3 and A21(N3) (3.2 Å). Atoms colored: DNA carbons (gray), oxygen (red), nitrogen (blue), phosphate (orange). MGB carbons: DAPI (purple), Hoechst (green), and netropsin (tan).

### Programming the lattice for site-specific positioning of MGBs

The structure of each molecule was solved at Pos1 or Pos2, yielding crystals with 50% of their minor grooves occupied. When the AATT sequence was positioned at BP, crystals containing full occupancy were acquired (Figure 2a-i). In addition, co-crystals prepared under identical conditions, but not bearing the binding sequence, yielded no discernible electron density at either MG. Each 4x5 Pos1 structure crystallized with *P*3_2_21 symmetry (as anticipated); however, Pos2 and BP crystallized with *P*3_2_ symmetry resulting in lattices with solvent channel volumes of ∼24 nm^3^ and ∼639 nm^3^, respectively.^55^ This dramatic difference was not entirely surprising given that subtle changes in junction angles or with downstream “flanking” sequence changes can alter symmetry,^55^ and although placement at Pos1 does not perturb the lattice, it appears that binding in Pos2 or BP influences the topology, causing the lattice to re-configure. Aside from the BP Hoechst structure (*R*3) all 4x6 crystals yielded *P*3_2_ lattices (Supplementary Table 4). The ability to position two separate guests within a single crystal will enable a range of applications where two different molecules need to be placed in close proximity, such as energy transfer or catalytic cascades.

### Electrostatic coordination of MGBs in the lattices

The most prominent feature of DAPI structures bound to AT rich MGs is the bifurcated hydrogen bonds the indole nitrogen forms with neighboring bases, with these characteristic contacts to T6(O2) and T14(O2) at Pos1, or T6(O2) and T19(O2) at Pos2 (Figure 3a-c) with additional interactions with the amidine moiety on each end of the molecule. However, although the 1D30 model was bound in a similar manner to the Pos2 structures (Figures 3a-c & Supplementary Figures 12a-b, 13a & 14a, we did observe that the binding mode of the molecule reverses when placed at Pos1 in both motifs. Nevertheless, the interactions with the floor of the MG remain the same, and the models and polar contacts are in good agreement with 1D30 (RMSDs=0.128-1.404; Supplementary Figures 13a, 14a & and 15).

Much like the previously reported versions of the Hoechst 33342, the coordination scheme of the 4x5 and 4x6 models is straightforward; however, the binding mode in 129D orients the ethoxyphenyl group on the 5’ end of the binding sequence (Supplementary Figure 12b), and this reverses with the piperazinyl leading at the 5’ in each of our models (Figure 3d-f). This change has no effect on the structure, where both benzimidazole moieties still provide the sole imidazole nitrogen groups that form three-centered hydrogen bonds with T6(O2) and T14(O2), and the other to A5(N3) and T14(O2) in the Pos1 structures. The similar binding scheme was observed in the Pos2 structures, as well as each of the 4x6 versions (Figure 3d-f & Supplementary Figure 12c-d). The details of the piperazinyl and ethoxyphenyl groups were also well justified with the electron density maps (Supplementary Figures 6-8d-f), and overall structures and electrostatic interactions track well with 129D (RMSDs=0.299-0.558; Supplementary Figures 13b, 14b & 16).

Netropsin is unique from DAPI and Hoechst in that it contains a repeating series of amide bonds that make the coordination schemes more complicated, but still predictable. In the 6BNA structure, guanidinium binds at the 5’ end, with the more flexible propylamidinium group on the other (Supplementary Figure 13c & 14c). However, we observe the opposite orientation can also be true even when the binding site is positioned in the same minor groove. This effect occurred in both motifs where netropsin binds at Pos2, where the guanidinium moiety binds at the 5’ end of the AATT sequence, but at Pos2 in the BP crystal, the propylamidinium appears at the 5’ end. Nevertheless, the amide-AATT interactions are unperturbed because the placement and proximity of the amides remains the same. For example, in either mode, the main amide hydrogen bonds to T4 and A18, T5 and T19, and the T20-amide contacts appear in each Pos2 structure, along with a host of other interactions on either end of the molecule (Figure 3g-i). This phenomenon also holds true in the 4x6 structures (Supplementary Figure 12e-g). In addition, we did observe that when the molecule is reversed, there is one additional amide interaction that occurs at the 3’ end of Chain A. We also note that each of the five-membered pyrrole rings are not coplanar, but rather run parallel to the walls of the MG. This is evident, along with the intervening interactions, in each of the resulting electron density maps, and is consistent with 6BNA (RMSDs=0.146-0.962; Figures 2g-i; Supplementary Figure 16).

In instances where there were apparent differences in the binding mode of Hoechst 33342 and netropsin compared to their previously described structures, it is not unreasonable to expect that this could be the case here. First, the context of the solvation sphere is non-trivial,^56^ and could lead to the preference for binding in one mode or another; however, at ∼3 Å it is not possible to realiably describe the hydration shell. Second, this effect could also be dictated by pH and salt concentration differences in the crystallization mother liquor itself. Lastly, these effects could also lead to slight perturbations of the width of the minor groove due to expansion or contraction of the unit cell which could be a determinant of the mode of interaction of minor groove binding molecules.^57^

### Crystal structure of ImPyPy-β-Dp (IPP) in the 4x5 lattice

We produced the IPP molecules using a similar synthetic route previously reported,^58^ but with slightly longer tails than those in the original NMR structure of (ImPyPy)_2_-TGTCA.^39,46,47^ Although this structure was described, we were unable to locate the coordinates in any of the common servers, and to our knowledge, the crystal structure of the (ImPyPy)_2_-TGTCA has not been solved. Nevertheless, the anticipated structure was expected to adhere to the design rules followed by previously described crystal structures of linear imidazole-pyrrole molecules of varying lengths bound to unique DNA binding sequences (e.g. PDB IDs: 1CVY, 1CVX, and 407D) with resolutions within the 2.15-2.27 Å resolution range.^59,60^

IPP was crystallized at Pos2 in the 4x5 system with *P*3_2_ symmetry, and yielded electron density that enabled model building of the molecule at 3.05 Å resolution Supplementary Figure 18). Although the initial maps did not show definitive separation for each IPP monomer, and the extended tails were not resolved, polder maps provided good evidence to corroborate the reliability of the built model, including the tail positions (Supplementary Figure 18a-c). The average c-axis of the single binder crystals was ∼60 Å; however, when the anti-parallel dimer “sandwiches” into the MG it expands to accommodate the molecule, causing the unit cell c-axis to extend to 62.2 Å. This slight reconfiguration of the cell is required to accommodate binding, and this crystallographic metric consistently indicated the presence of bound IPP. Interestingly, unlike in other scenarios where slight geometric perturbances alter the crystal symmetry,^55^ the c-axis expansion does not disrupt the lattice.

As discussed above, deliberate efforts had been made towards the identification of pyrrole-imidazole polyamides with affinities and specificities comparable to DNA-binding proteins. In the NMR structure, when the (ImPyPy)_2_-TGTCA antiparallel dimer orients an imidazole on one side stacked to a pyrrole carboxamide on the opposing molecule, it strictly distinguishes a G•C pair; however, when the converse is true, only a C•G pair is recognized. Py-Py partners do not interact with either G•C or C•G pairs, but cannot discriminate between A•T/T•A base pairs.^39,46,47^

The polar contacts of the IPP molecule bound at Pos2 alone (Supplementary Figure 18d), and in the crystal containing both the netropsin and the IPP at Pos2 (Figure 4a; Supplementary Figure 19) were examined in parallel, and were both consistent with the predicted main interactions with the MG sequence, with the only discrepancies arising in the flexible tails. Due to the quality of the electron density (Figure 4a-b; Supplementary Figure 20a-c) and much higher resolution (2.45 Å) in the relative regime of previous polyamide structures, a detailed description of the structural contacts and distances, from the netropsin:IPP crystal of the polyamide was possible (Figure 4b & Supplementary Figure 19). Specifically, the polyamide strictly adhered to the sequence design rules that had been originally designed for the bound homodimer. The structure revealed that both the Im/Py (G17•C7) and Py/Im (C19•G5) maintained their exclusive preference for G•C and C•G sequences respectively, and that the degenerate Py/Py pair contacts T18•A6, but did not interact with G•C/C•G pairs. Multiple other contacts with the tail regions were also observed (Figure 4c). The ability to properly build the model and map the molecular interactions, owes in no small part to the data quality and resolution, and the exquisite detail revealed in the resulting electron density (Figure 4a; Supplementary Figure 20).

**Figure 4.**
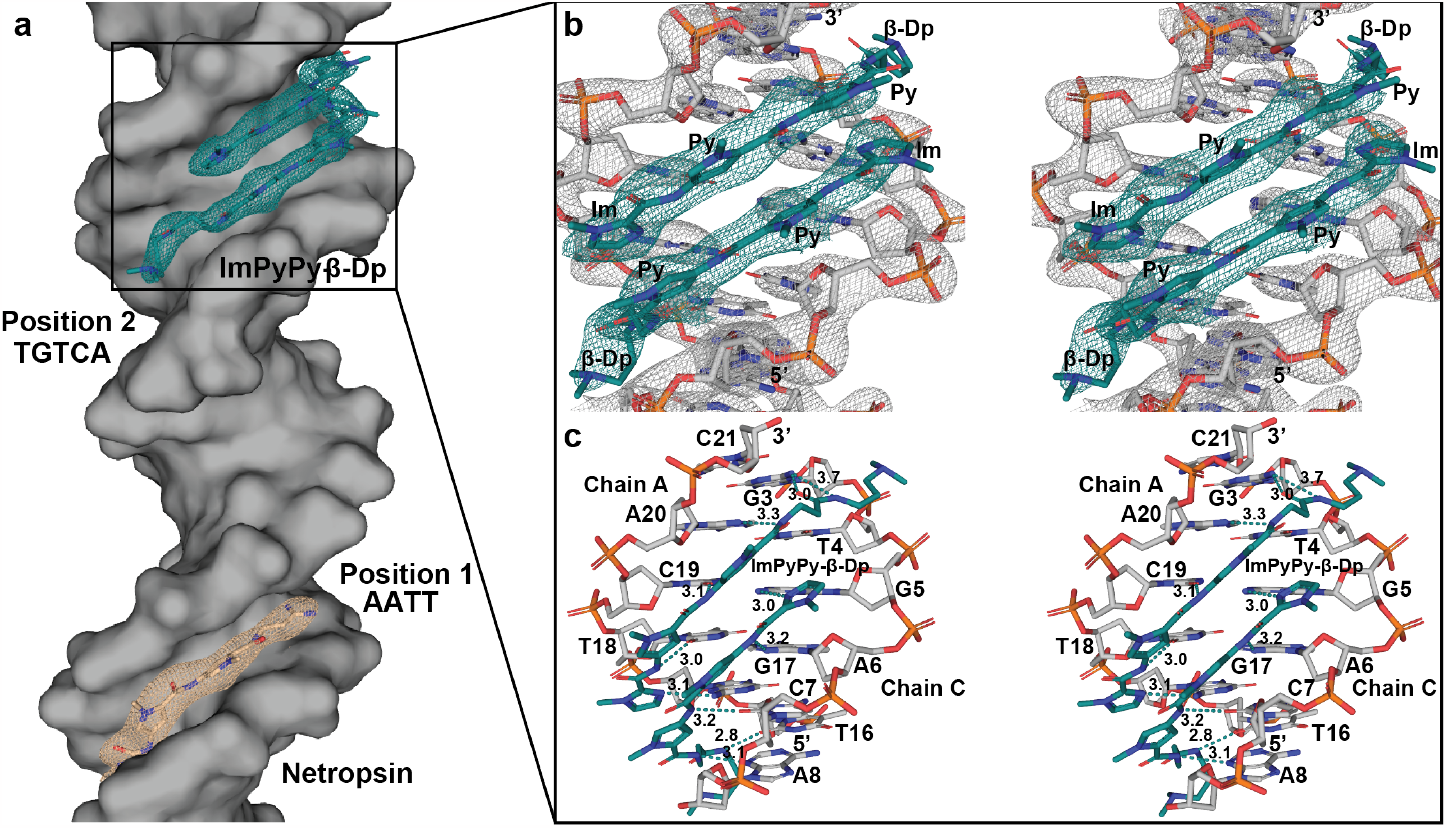
Crystal structure and electron density for bound IPP and mapped molecular contacts. (a) The structure of a single duplex from a co-crystal containing netropsin (tan) and IPP (teal). The DNA contained an AATT sequence at minor groove Pos1 which is recognized by the bound netropsin molecule and TGTCA at Pos2 which is the cognate sequence required for the binding of the IPP polyamide. The 2*F*_*o*_ *– F*_*c*_ electron density (tan) for the netropsin molecule is contoured at σ = 1.0, and σ = 1.5 (teal) for the IPP; (b) Stereoscopic view of the 2*F*_*o*_ *– F*_*c*_ electron density contoured at σ = 2.5 (gray) of the DNA containing the TGTCA minor groove binding sequence. The map was contoured such that the phosphate backbone, base stacking, and map/model quality is evident. The 2*F*_*o*_ *– F*_*c*_ electron density contoured at σ = 1.5 (teal) of the bound IPP molecules is also shown. The anti-parallel orientation of each monomer and the constituent stacked pyrrole and imidazole pairs (labeled) are well accounted for in the electron density of the 2.45 Å model. The long β-Dp tails were expected to have a fair degree of flexibility, and thus complete density coverage was not achievable at the 3’ end of the top monomer; however, polder maps demonstrated additional density to account for the reliability of the built model (Supplementary Figure 19). (c) Stereoscopic view of the mapped polar contacts between the anti-parallel IPP dimer and the floor of the minor groove. The 5’ end of the TGTCA (16-20) in chain A begins in the lower right. Polar contacts are indicated with teal dashes. Electrostatic interactions were defined between atoms of distances of 2.5 – 3.7 Å. The G3•C21 pair at the top is included in the sequence because the β-Dp tail forms two additional contacts with G3 on the 5’ end of chain C. All modeled contacts are accounted for between each of the pyrrole and imidazole stacking combinations with the designed recognition sequence in the minor groove. Atom coloring for both panels are as follows: DNA carbons (gray) and IPP carbons (teal), oxygen (red), nitrogen (blue), phosphate (orange).

### Symmetry dictates lattice presentation of MGBs

Along with its smaller MGB counterparts in a *P*3_2_ lattice, the IPP Pos2 structure is best viewed as a cross-section of four layers with a “front” and “reverse” face (Figure 5a; Supplementary Figures 21-25a), as well as for the 4x6 *R*3 Hoechst BP crystals, which also displays the molecule in the same fashion (Supplementary Figure 25b). When viewed along the front face of the crystal, all molecules face forward, and the difference in occupancy between the single positions and BP crystals is apparent. Further, when the crystal is oriented 180° from the front, no molecules are visible in the MGs.

**Figure 5.**
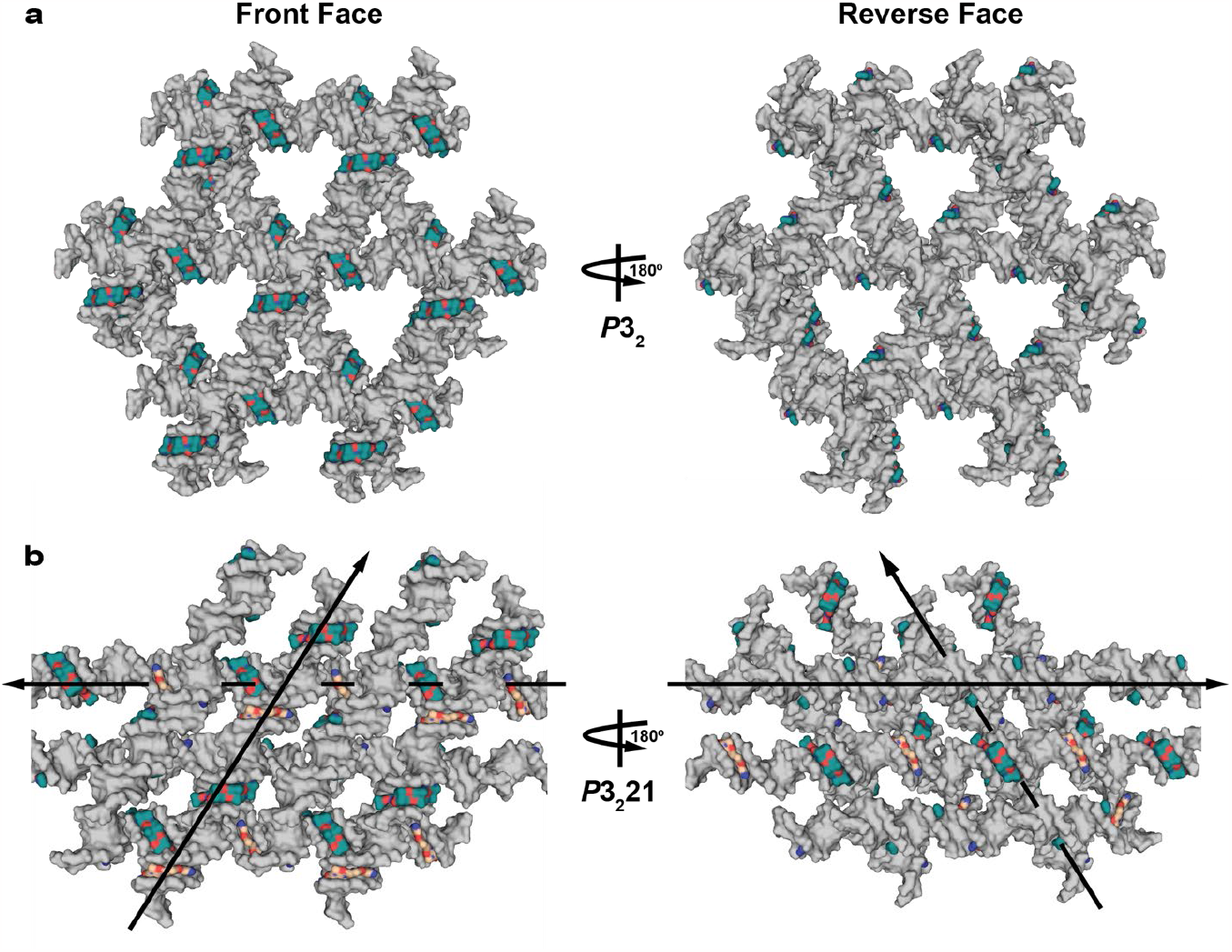
Symmetry dependent lattice packing and orientation of IPP in the 4x5 crystals. (a) *P*3_2_ lattice of the 4x5 Pos2 crystals containing a surface representation of the bound IPP dimer. The molecules bind at alternating minor grooves which contain the TGTCA binding sequence, and present only on the front face of the crystal in each layer, whereas when the crystal is oriented 180° from the front side, no molecules are visible on the reverse face. (b) *P*3_2_21 lattice of the 4x5 crystal with netropsin bound at Pos1 and IPP bound at Pos2. A cross-section of two layers demonstrating that the molecules on a front facing crystal present in both positions on alternating duplexes in each layer. Arrows are used to indicate which layer faces forward, and that each layer stacks across the major grooves between each binder on the opposite layer. The same is also true upon 180° degree rotation to the opposing face. The surface of the IPP (teal carbons) and netropsin (tan carbons) molecules have been surface rendered, with oxygen (red) and nitrogen (blue) also represented.

The difference between the presentation of the small MGBs in the 4x5 Pos1 crystals and of IPP and netropsin in the mixed lattice is dramatic, due to the *P*3_2_21 symmetry. When the crystal is observed as a two-layer cross-section, the MGs in each constituent layer contain a bound target at repeating intervals, but only on alternating linear helices within each layer. The molecules oriented on the backside of each linear duplex is observed when oriented 180° from the “top” layer, and is consistent with the same presentation of the molecules in the 4x5 Pos1 structures of the small MGBs (Figure 5b and Supplementary Figure 26).

The ability to orient bound molecules based upon crystal symmetry provides an attractive route towards engineering crystals that not only sequence-specifically position guests, but also with the ability to spatially arrange these elements in a manner dictated by the symmetry of the scaffold itself. Unlike previously reported scenarios where MG intercalating drug molecules are studied crystallographically,^61^ rigid, self-assembled DNA lattices that are designed to “automatically” crystallize, have not been used, and could serve as highly modular frameworks as hosts for a large variety of intercalators. This becomes apparent when the guest molecules within the entire lattice are visualized in 3D space (see videos in Section SX). We envision that modifying functional groups (such as enzymes, small molecule catalysts, or light-harvesting dyes) with the MG binders could provide a facile and modular way to scaffold them in the crystal with defined distances and relative orientations. The use of two different binders in a single crystal is particularly attractive because it enables the strategic placement of enzyme pairs, nanoparticles, or FRET dyes, for example, to create catalytic cascades or photonic/excitonic materials.

## CONCLUSION

We solved the structures of a panel of three unique MGBs, bound to their consensus binding sequence (AATT) in self-assembled DNA crystals. We further demonstrated that despite differences in symmetry, the scaffolds could be used to effectively orient the molecules with high resolution. It was also possible to control the occupancy of the MGB by tuning the position(s) of the sequence within the MGs, and thereby the fraction of sites available for binding. With this capability, we solved the structure of IPP bound to TGTCA at Pos2, and with netropsin at Pos1 to demonstrate that multiple molecules could be tethered within these assemblies, and solved the 2.45 Å IPP structure with all of its molecular contacts previously reported in an NMR structure remaining intact, highlighting its designed sequence selectivity.

DNA crystal motifs provide a highly versatile environment towards studying a variety of biological molecules, and MGBs could effectively provide an exciting new route towards stably anchoring conjugated guests with high affinity to prescribed positions in the DNA lattice for characterization. This methodology would be an attractive strategy for the placement of small molecules, nanoparticles, or proteins, e.g. by chemically linking them to the MG-binding small molecules. The findings can open new avenues towards the use of DNA motifs for structural biology and as functional scaffolds, and demonstrates that structural DNA nanotechnology continues to provide great promise for probing the structural and functional properties of biological phenomena.

## MATERIALS & METHODS

### Synthesis and characterization of ImPyPy-β-Dp

The IPP minor groove binder was obtained via solid-phase synthesis through standard Fmoc-protected monomers, prepared in the laboratory using an adaption of a previously described protocol.^58^ Peptide synthesis was performed in a peptide synthesis reaction vessel (ChemGlass, CG-1860) with each reaction step carried out on a Burrel Scientific wrist action shaker. The molecule was synthesized on Fmoc-β-Ala-Wang (APPTEC) resin at a 0.1 mmol scale. Deprotection of the Fmoc protecting group was accomplished by shaking the resin in 10 mL of 20% piperidine in NMP for 20 minutes. A 5-fold excess of each monomer and HBTU were mixed in DMF with 1 mL of pure DIPEA. This mixture was allowed to react for 5 minutes to pre-activate the carboxylic acid and was then added to the shaking peptide resin for 1 hour. After synthesis of the molecule was complete, it was cleaved from the resin through aminolysis by incubating the resin in 4 mL of dimethylaminopropylamine at 55°C for 18 hours, after which the cleavage solution was mixed with 16 mL of 50:50 water:acetonitrile. The crude product was then purified on a Waters HPLC using a C18 Phenomenex column. Purified fractions were analyzed on an AB SCIEX 4800 MALDI TOF/TOF using α-cyano-4-hydroxycinnamic acid matrix to confirm successful synthesis and purity. The pooled peak fractions containing the pure molecule were lyophilized and stored at -20°C.

### Crystallization and sample preparation

Netropsin dihydrochloride was purchased from Santa Cruz Biotechnologies. Hoechst 33342 trihydrochloride trihydrate and 4′,6-diamidino-2-phenylindole (DAPI) were purchased from Thermo Fisher Scientific. Stock solutions of MGBs were resuspended in water to a final concentration of 5 mg/mL. All oligonucleotides were purchased from Integrated DNA Technologies (Coralville, IA), purified by 14% denaturing polyacrylamide gel electrophoresis (PAGE), and subsequently resuspended to a final concentration of 300 μM in Nanopure water. Stock solutions for all three component DNA strands and MGBs were prepared at a 30:120:120:X μM ratio (S1:S2:S3:MGB) with “X” being variable depending on the screening condition that yielded the respective crystal structure. Sparse matrix screening was performed using the sitting drop vapor diffusion method in Cryschem plates (Hampton Research) with an adaptation of a discontinued DNA crystallization screen of 48 conditions (Sigma Aldrich). Crystallization was carried out with 500 µL of reservoir buffer and a total drop volume of 6 µL with a 2:1 ratio of DNA/MGB stock to buffer. The crystal trays were heated in a chilling incubator (Torrey Pines Scientific; Carlsbad, CA) to 60°C for 1 hour and then cooled to 25°C using a linear gradient at a rate of 0.3°C/hr. Crystal “hits” from the initial screening were then optimized by probing conditions such as MGB and DNA concentration, salt or solvent concentration, pH, surfactant concentration as well as the rate of the temperature gradient to enhance crystal quality. The resulting crystals were imaged using a light microscope (Supplementary Figures 2, 3 & 5), with the corresponding crystallization conditions in (Supplementary Tables 2, 3 & 5). Sample harvesting was performed using cryo-loops (Hampton Research) by placing the crystals into a drop containing artificial mother liquor supplemented with 30% glycerol, and then cryo-cooled by immediate submersion into a liquid nitrogen bath.

### Data Collection, Processing and Structure Solution

All diffraction data were acquired in a nitrogen cold stream (100 K) at multiple light sources, as indicated in (Supplementary Tables 6-9). Data were processed using HKL2000.^62^ Phases for the resulting structures were derived via molecular replacement with either Phaser^63^ or Molrep^64^ from the CCP4^65^ suite of programs using either 5KEK^21^ (4x5) or 5VY6^22^ (4x6) as the search models for the corresponding motif that was used, and yielded positive *F*_*o*_ *– F*_*c*_ difference Fourier density corresponding to each respective MGB. The initial solutions were then treated as a single rigid body, and manual model building commenced in Coot^66^. Multiple iterative rounds of model building and refinement was performed using restrained refinement in REFMAC^67^ from CCP4, in tandem with XYZ coordinate, real space, occupancy, and B-factor calculation refinements using phenix.refine from the Phenix software package.^68^ Water and other ion positions resulting from the mother liquor were interpreted based upon 2*F*_*o*_ *– F*_*c*_ peaks ≥ σ = 1.0, and subsequently refined. Rigid body refinement was then performed to conclude refinement of the non-ligand bound structure once acceptable R-factors had been achieved using an Rfree set containing 5–10% of the unique reflections. Known ligand coordinates (DAP, HT1, and NT) for DAPI, Hoechst 33342, and netropsin, respectively were automatically retrieved in Phenix, and the IPP structure was drawn in Chemdraw. Quantum calculations to provide an accurate definition file for each molecule was performed using elBOW.^69^ Subsequent to completion of model building and refinement, polder^52^ omit maps were calculated via the omission of the individual molecules to allow for the exclusion of bulk solvent in the omitted region to further substantiate that the model was built correctly, and in good agreement with the 2*F*_*o*_ *– F*_*c*_ electron density maps.

## ASSOCIATED CONTENT

### Supporting Information

The contents of the supplementary information include the topological schematics and sequences of the crystal designs, the crystallographic data collection and refinement statistics, crystal images, 2*F*_*o*_ *– F*_*c*_ and polder electron density maps with superpositions demonstrating map agreement at all positions in both the 4x5 and 4x6 systems, structure and electrostatic coordination of DAPI, Hoechst, and netropsin in the 4x6 motif, structure and electrostatic interactions and alignment statistics of 1D30, 129D, and 6BNA with 2D schematics of mapped interactions for DAPI, Hoechst, netropsin in these structures, 2D schematic of mapped IPP-TGTCA interactions in the 4x5 Pos2 crystal with resulting electron density, and the lattice packing and orientation views of all MGBs in the 4x5 and 4x6 systems.

### Accession codes

8TB3, 8TB4, 8T7X, 8TA9, 8TDT, 8TB8, 8TBD, 8T80, 8TAQ, 8TAM, 8TBO, 8T7B, 8TC2, 8TAJ, 8TAP, 8TA8, 8TC4, 8TC6

The Supporting Information is available free of charge on the ACS Publications website.

## Author Contributions

All authors have given approval to the final version of the manuscript.

## ACKNOWLEDGMENT

Results shown in this report are derived from work performed at the Light Source (ALS), Argonne Photon Source (APS), and the National Synchrotron Light Source II (NSLS-II). We gratefully acknowledge beamline 5.0.2 of the ALS, a DOE Office of Science User Facility under Contract No. DE-AC02-05CH11231, which is supported in part by the ALS-ENABLE program funded by the National Institutes of Health, National Institute of General Medical Sciences, grant P30 GM124169-01, and for research performed at the Structural Biology Center which is funded by the U.S. Department of Energy, Office of Biological and Environmental Research and operated for the DOE Office of Science at the APS by Argonne National Laboratory under Contract No. DE-AC02-06CH11357. Additionally, this research used resources AMX (17-ID) and FMX (17-BM) of the NSLS-II, a U.S. Department of Energy (DOE) Office of Science User Facility operated for the DOE Office of Science by Brookhaven National Laboratory under Contract No. DE-SC0012704. The Center for BioMolecular Structure (CBMS) is primarily supported by the National Institutes of Health, National Institute of General Medical Sciences (NIGMS) through a Center Core P30 Grant (P30GM133893), and by the DOE Office of Biological and Environmental Research (KP1607011). N.S. acknowledges that research reported in this publication was supported by The National Institute of General Medical Sciences of the National Institutes of Health under grant number DP2GM132931. N.S. and H.Y. gratefully acknowledge support from the National Science Foundation Division of Materials Research (NSF2004250). H.Y. additionally acknowledges financial support from Arizona State University.

## ABBREVIATIONS

DAPI: 4’,6-diamidino2-phenylindole;
IPP: imidazole-pyrrolepyrrole;
MG: minor groove;
MGB: minor groove binder;
Pos1: position 1;
Pos2: position 2;
S1, BP: both positions; strand 1
S2: strand 2;
S3: strand 3

