## Supplementary Information for "Site-specific arrangement and structure determination of minor groove binding molecules in self-assembled three-dimensional DNA crystals"

### Table of Contents

**Supplementary Figure 1.** Topological schematics of the 4x5 and 4x6 crystal motifs 4
**Supplementary Table 1.** DNA sequences used for each constituent oligonucleotide 5
**Supplementary Table 2.** Components for the 48 buffer conditions used for crystallization screening 6
**Supplementary Figure 2.** Representative bright field images of the 4x5 MGB co-crystals 7
**Supplementary Table 3.** The buffer conditions used for the corresponding bright field images 7
**Supplementary Figure 3.** Representative bright field images of the 4x6 MGB co-crystals 8
**Supplementary Table 4.** Buffer conditions used for the corresponding bright field images 8
**Supplementary Figure 4.** Topological schematic of the 4x5 motif for binding of netropsin and IPP at unique positions 9
**Supplementary Figure 5**. Representative bright field images of the 4x5 IPP Pos2 and IPP Pos2 + netropsin Pos1 co-crystals 10
**Supplementary Table 5.** Crystallization buffers corresponding to the bright field images 10
**Supplementary Table 6.** Data collection and refinement statistics for DAPI containing crystal 11
**Supplementary Table 7.** Data collection and refinement statistics for Hoechst containing crystals 12
**Supplementary Table 8.** Data collection and refinement statistics for netropsin containing crystals 13
**Supplementary Table 9.** Data collection and refinement statistics for IPP and IPP + netropsin crystals 14
**Supplementary Figure 6.** Electron density maps and structures of MGBs bound at Pos1 in the 4x5 motif 15
**Supplementary Figure 7.** Electron density maps and structures of MGBs bound at Pos2 in the 4x5 motif 16
**Supplementary Figure 8.** Electron density maps and structures of MGBs bound at Pos2 in the 4x5 motif 17
**Supplementary Figure 9**. Electron density maps and structures of DAPI at Pos1 and Pos2 in the 4x6 motif 18
**Supplementary Figure 10.** Electron density maps and structures of Hoechst molecules bound at Pos2 and BP in the 4x6 motif 19
**Supplementary Figure 11.** Electron density maps and structures of netropsin bound at Pos1, Pos2 and BP in the 4x6 motif 20
**Supplementary Figure 12.** Structure and electrostatic coordination of DAPI, Hoechst, and netropsin to the AATT minor groove in the 4x6 lattices 21
**Supplementary Table 10.** Structure alignment of DAPI, Hoechst, and netropsin (1D30, 129D, and 6BNA) 22
**Supplementary Figure 13**. Reference molecule superpositions of MGB structures to original crystal structures 22
**Supplementary Figure 14**. Structure and electrostatic coordination of DAPI (1D30), Hoechst (129D), and netropsin (6BNA) to the AATT minor groove in the original structures 23
**Supplementary Figure 15.** 2D schematic of the DAPI-AATT polar contacts discovered in PDB 1D30 24
**Supplementary Figure 16.** 2D schematic of the Hoechst-AATT polar contacts discovered in PDB 129D 25
**Supplementary Figure 17.** 2D schematic of the netropsin-AATT polar contacts discovered in PDB 6BNA 26
**Supplementary Figure 18.** Electron density and polar contacts of the IPP Pos2 structure 27
**Supplementary Figure 19.** 2D schematic of the IPP-TGTCA polar contacts in the 4x5 IPP + netropsin co-crystal 28
**Supplementary Figure 20**. Electron density containing the structures of IPP and netropsin in the 4x5 29

**Supplementary Figure 21.** 4x5 Pos2 crystal lattice packing and molecule orientation **30**
**Supplementary Figure 22.** 4x5 crystal lattice packing and orientation at both positions **31**
**Supplementary Figure 23.** 4x6 crystal lattice packing and orientation of DAPI molecules **32**
**Supplementary Figure 24.** 4x6 crystal lattice packing and orientation of netropsin molecules **33**
**Supplementary Figure 25.** 4x6 crystal lattice packing and orientation of Hoechst molecules **34**
**Supplementary Figure 26.** 4x5 Pos1 crystal lattice packing and orientation **35**

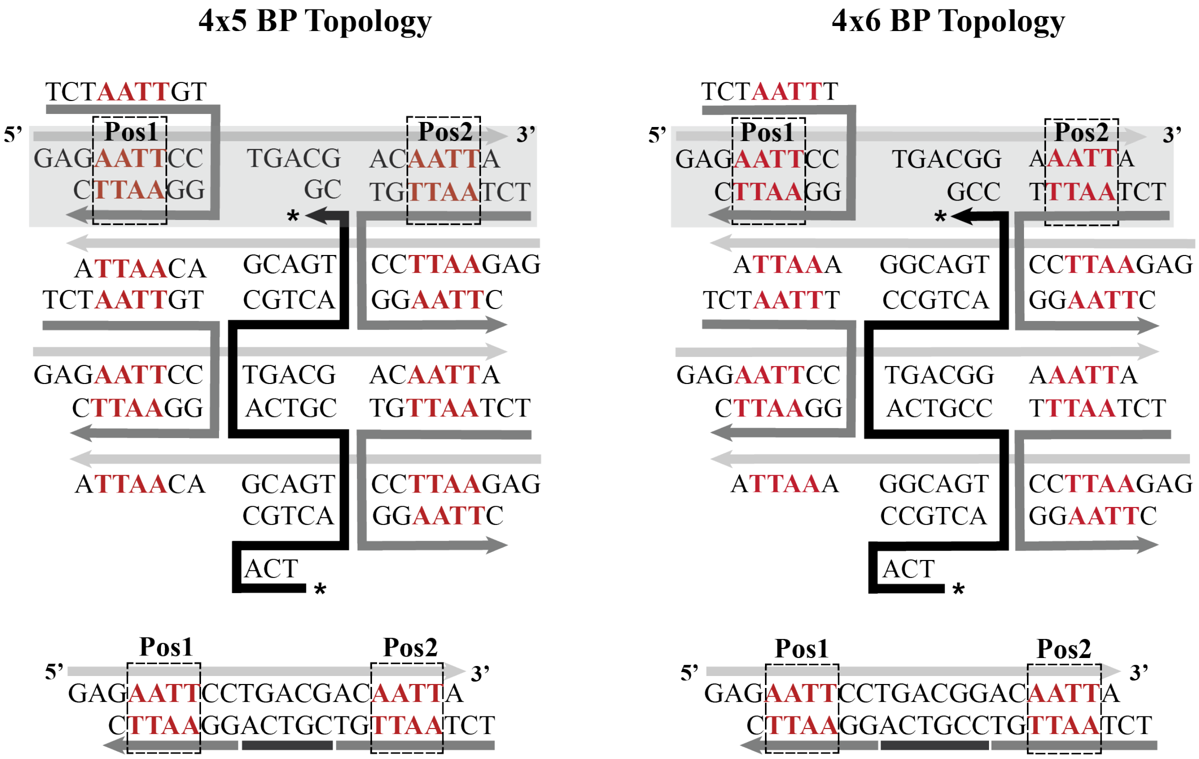

**Supplementary Figure 1. Topological schematics of the 4x5 and 4x6 crystal motifs.** 2D topological representations of each crystal motif. The central "building block" of each motif is made up of three component oligonucleotides. S1 (black) contains four repeats of either 5 (left) or 6 (right) bases that tether four 21-bp duplexes (two full helical turns) through a series of four-arm Holliday junction crossovers; S2 (light gray) is a linear 21-base oligonucleotide that does not participate in any junction crossover, and comprises one side of each duplex; and S3 (dark gray) forms the second half of each Holliday junction, and serves as a connection between two arms between adjacent layers. The asterisks indicate where the 5' end of S1 hybridizes to its complement in the fourth duplex. The gray box highlights a duplex within the asymmetric unit of the crystal from a single layer (bottom left and right). Each duplex is tailed by two base complementary "sticky ends" which hybridize to connect each motif block within the crystal lattice. Each 21-bp duplex contains two minor grooves (Pos1 and Pos2). Three versions of each motif were created with the AATT binding sequence (red) at either Pos1 or Pos2, or at both positions. In the 4x5 crystals, the Pos1 and Pos2 binding sequences were placed on the 5' and 3' ends of S2, one base away from the sticky end and two bases from the junction, to ensure that each site was unperturbed upon co-crystallization. The 4x6 motif was fashioned in a similar manner.

**Supplementary Table 1. DNA sequences used for each constituent oligonucleotide.** The DAPI, Hoechst, and netropsin AATT binding sequences are highlighted in orange, and the IPP-TGTCA is shown in blue.

| Netropsin, DAPI, Hoechst | | |
|---|---|---|
| **Motif** | **Strand** | **Sequence** |
| **4x5 Pos. 1** | S1 | TCACGTCACGTCACGTCACG |
| | S2 | GACAATTGCTGACGACACTCA |
| | S3 | TCTGAGTGTGCAATTG |
| **4x5 Pos. 2** | S1 | TCACGTCACGTCACGTCACG |
| | S2 | GAGCAGACCTGACGACAATTA |
| | S3 | TCTAATTGTGGTCTGC |
| **4x5 Both Pos.** | S1 | TCACGTCACGTCACGTCACG |
| | S2 | GAGAATTCCTGACGACAATTA |
| | S3 | TCTAATTGTGGAATTC |
| **4x6 Pos. 1** | S1 | TCACCGTCACCGTCACCGTCACCG |
| | S2 | GAGAATTCCTGACGGAACTCA |
| | S3 | TCTGAGTTGGAATTC |
| **4x6 Pos. 2** | S1 | TCACCGTCACCGTCACCGTCACCG |
| | S2 | GAGCAGACCTGACGGAAATTA |
| | S3 | TCTAATTTGGTCTGC |
| **4x6 Both Pos.** | S1 | TCACCGTCACCGTCACCGTCACCG |
| | S2 | GAGAATTCCTGACGGAAATTA |
| | S3 | TCTAATTTGGAATTC |

| ImPyPy | | |
|---|---|---|
| **Motif** | **Strand** | **Sequence** |
| **4x5 Pos. 2** | S1 | TCACGTCACGTCACGTCACG |
| | S2 | GAGCAGACCTGACGATGTCAC |
| | S3 | TCGTGACATGGTCTGC |

| ImPyPy + Netropsin | | |
|---|---|---|
| **Motif** | **Strand** | **Sequence** |
| **4x5 Both Pos.** | S1 | TCACGTCACGTCACGTCACG |
| | S2 | GACAATTGCTGACGATGTCAC |
| | S3 | TCGTGACATGCAATTG |

### Supplementary Table 2. Components for the 48 buffer conditions used for crystallization screening

| | | | | | | |
|---|---|---|---|---|---|---|
| **1** | 0.05 M HEPES pH 7.5 | 80 mM $MgCl_2$ | 2.5 mM spermine | | | |
| **2** | 0.05 M Na cacodylate pH 6.0 | 18 mM $MgCl_2$ | 2.25 mM spermine | 1 mM $CuSO_4$ | 9% isopropanol | |
| **3** | 0.05 M Na cacodylate pH 6.5 | 18 mM $MgCl_2$ | 0.9 mM spermine | 1.8 mM $CoH_{18}N_6$ | 9% isopropanol | |
| **4** | 0.05 M Na cacodylate pH 6.5 | 18 mM $MgCl_2$ | 2.25 mM spermine | 9% isopropanol | | |
| **5** | 0.05 M Na cacodylate pH 7.0 | 18 mM $MgCl_2$ | 2.25 mM spermine | 0.9 mM $CoH_{18}N_6$ | 4.5% MPD | |
| **6** | 0.05 M Na cacodylate pH 6.5 | 36 mM $MgCl_2$ | 2.25 mM spermine | 5% PEG 400 | | |
| **7** | 0.05 M Na succinate pH 5.5 | 10 mM $MgCl_2$ | 2.0 mM $CoH_{18}N_6$ | 10% isopropanol | | |
| **8** | 0.05 M Na cacodylate pH 6.0 | 20 mM $MgCl_2$ | 1.0 mM spermine | 15% ethanol | | |
| **9** | 0.05 M Na cacodylate pH 7.0 | 20 mM $MgCl_2$ | 1.0 mM spermine | 1.0 mM $CoH_{18}N_6$ | 15% ethanol | |
| **10** | 0.05 M Na cacodylate pH 7.0 | 5 mM $MgCl_2$ | 1.0 mM spermine | 10% tert-butanol | | |
| **11** | 0.05 M Na cacodylate pH 7.0 | 30 mM $MgCl_2$ | 2.5 mM spermine | 5% PEG 400 | | |
| **12** | 0.05 M Na cacodylate pH 6.5 | 100 mM $MgCl_2$ | 2.0 mM $CoH_{18}N_6$ | 5% isopropanol | | |
| **13** | 0.05 M Tris pH 8.0 | 10 mM $MgCl_2$ | 1.0 mM $CoH_{18}N_6$ | 20% ethanol | | |
| **14** | 0.05 M HEPES pH 7.5 | 20 mM $MgCl_2$ | 1.0 mM spermine | 5% PEG 8000 | | |
| **15** | 0.05 M Na cacodylate pH 6.0 | 20 mM $MgCl_2$ | 2.5 mM spermine | 5% PEG 4000 | | |
| **16** | 0.05 M Na cacodylate pH 6.0 | 10 mM $MgCl_2$ | 2.5 mM spermine | 5 mM $CaCl_2$ | 10% isopropanol | |
| **17** | 0.05 M Na cacodylate pH 7.0 | 9 mM $MgCl_2$ | 2.25 mM spermine | 1.8 mM $CoH_{18}N_6$ | 0.9 mM spermidine | 5% PEG 400 |
| **18** | 0.05 M Na cacodylate pH 6.5 | 10 mM $MgCl_2$ | 2.5 mM spermine | 1 mM $CuSO_4$ | 10% isopropanol | |
| **19** | 0.05 M Na cacodylate pH 6.0 | 20 mM $MgCl_2$ | 1.0 mM spermine | 2 mM $CaCl_2$ | 10% MPD | |
| **20** | 0.05 M HEPES pH 7.5 | 15 mM $MgCl_2$ | 1.0 mM spermidine | 10% dioxan | | |
| **21** | 0.05 M Na cacodylate pH 6.0 | 15 mM $MgCl_2$ | 3.0 mM spermine | 10% PEG 400 | | |
| **22** | 0.05 M Na cacodylate pH 6.5 | 2.5 mM spermine | 18 mM $CaCl_2$ | 9% isopropanol | | |
| **23** | 0.05 M Na cacodylate pH 6.5 | 2.0 mM spermine | 1.0 mM $CoH_{18}N_6$ | 80 mM $CaCl_2$ | | |
| **24** | 0.05 M Na cacodylate pH 6.5 | 5 mM $MgCl_2$ | 2.5 mM $CoH_{18}N_6$ | | | |
| **25** | 0.05 M Na cacodylate pH 6.5 | 30 mM $MgCl_2$ | 1.0 mM spermine | 1.3 M $Li_2SO_4$ | | |
| **26** | 0.05 M Na cacodylate pH 6.0 | 200 mM $Ca(CH_3COO)_2$ | 5% isopropanol | | | |
| **27** | 0.05 M Na cacodylate pH 6.5 | 100 mM MgCl2 | 1.0 mM $CoH_{18}N_6$ | 10% ethanol | | |
| **28** | 0.05 M Na cacodylate pH 6.0 | 10 mM $MgCl_2$ | 2.5 mM spermidine | 2.5 M NaCl | | |
| **29** | 0.05 M Na cacodylate pH 6.5 | 10 mM $MgCl_2$ | 200 mM sodium citrate | 5% isopropanol | | |
| **30** | 0.05 M Na cacodylate pH 6.5 | 15 mM $MgCl_2$ | 10 mM spermine | 2.0 M $Li_2SO_4$ | | |
| **31** | 0.05 M Na cacodylate pH 6.5 | 20 mM $MgCl_2$ | 1.0 mM spermine | 2.0 M $(NH_4)_2SO_4$ | | |
| **32** | 0.05 M Na cacodylate pH 6.5 | 10 mM $MgCl_2$ | 1.5 mM spermine | 3.0 M $(NH_4)_2SO_4$ | | |
| **33** | 0.05 M HEPES pH 7.5 | 15 mM $MgCl_2$ | 1.0 mM spermine | 1.0 M $(NH_4)_2SO_4$ | | |
| **34** | 0.05 M Na cacodylate pH 6.0 | 200 mM $Ca(CH_3COO)_2$ | 2.5 M NaCl | | | |
| **35** | 0.05 M Na cacodylate pH 6.0 | 200 mM $Ca(CH_3COO)_2$ | 1.0 mM $CoH_{18}N_6$ | 2.0 M LiCl | | |
| **36** | 0.05 M Na cacodylate pH 6.5 | 15 mM $MgCl_2$ | 5.0 mM spermidine | 1.0 mM $CoH_{18}N_6$ | 2.0 M NaCl | |
| **37** | 0.05 M Na cacodylate pH 6.5 | 200 mM $MgCl_2$ | 100 mM NaCl | 20% PEG 1000 | | |
| **38** | 0.05 M Tris pH 7.5 | 50 mM $MgCl_2$ | 1.0 M sodium tartarate | | | |
| **39** | 0.05 M Tris pH 7.5 | 200 mM $MgCl_2$ | 2.5 M NaCl | | | |
| **40** | 0.05 M Na cacodylate pH 6.0 | 200 mM $MgCl_2$ | 2.5 M KCl | | | |
| **41** | 0.05 M Tris pH 8.0 | 200 mM $MgCl_2$ | 15% ethanol | | | |
| **42** | 0.05 M Na cacodylate pH 6.0 | 15 mM $MgCl_2$ | 5.0 mM spermidine | 2.0 M $Li_2SO_4$ | | |
| **43** | 0.05 M Na cacodylate pH 6.0 | 20 mM $Mg(CH_3COO)_2$ | 0.5 mM spermine | 100 mM NaCl | 25% MPD | |
| **44** | 0.05 M Na succinate pH 5.5 | 20 mM $MgCl_2$ | 0.5 mM spermine | 3.0 M $(NH_4)_2SO_4$ | | |
| **45** | 0.05 M Na cacodylate pH 6.5 | 5.0 mM $CoH_{18}N_6$ | 2.5 M KCl | | | |
| **46** | 0.05 M Na cacodylate pH 6.5 | 50 mM $MgCl_2$ | 2.0 mM $CoH_{18}N_6$ | 1.5 M $Li_2SO_4$ | | |
| **47** | 0.05 M Na cacodylate pH 6.5 | 1.0 mM spermine | 2.0 mM $CoH_{18}N_6$ | 30 mM $CaCl_2$ | 2.0 M LiCl | |
| **48** | 0.05 M Na cacodylate pH 6.5 | 10 mM $MgCl_2$ | 50 mM spermine | | | |

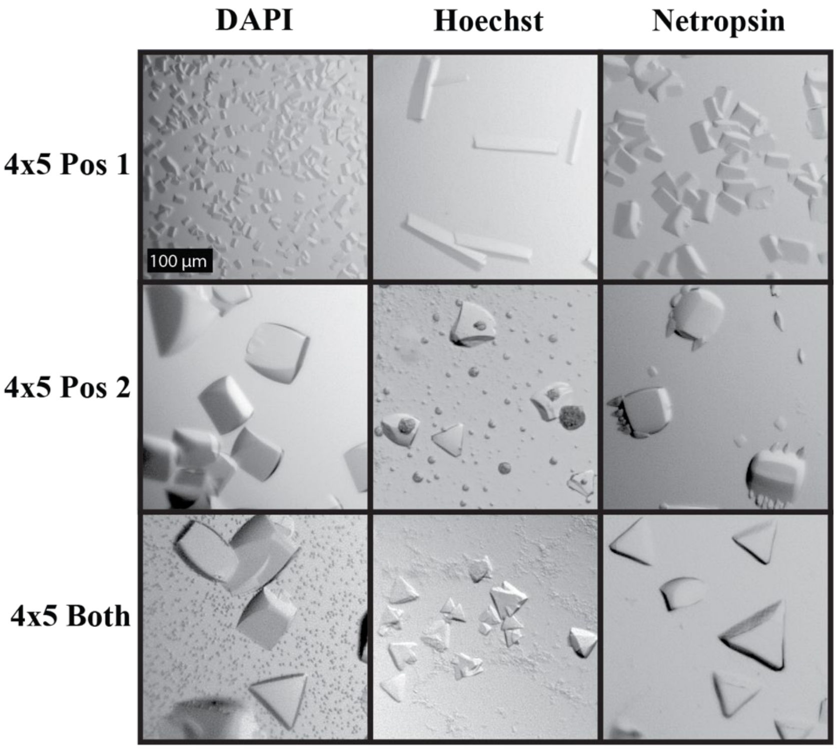

**Supplementary Figure 2. Representative bright field images of the 4x5 MGB co-crystals.** The buffers that correspond to each image are indicated in Supplementary Table 3.

**Supplementary Table 3.** The buffer conditions used for the corresponding bright field images.

| 4x5 Motif | | | | | | | |
|---|---|---|---|---|---|---|---|
| **Netropsin** | | | | | | | |
| **Motif** | **Binding Ratio** | **Buffer** | **Buffer Components** | | | | |
| Pos. 1 | 2X | 5 | 0.05 M Na Cacodylate pH 7.0 | 18 mM $MgCl_2$ | 2.25 mM spermine | 0.9 mM $CoH_{18}N_6$ | 4.5% MPD |
| Pos. 2 | 6X | 31 | 0.05 M Na Cacodylate pH 6.5 | 20 mM $MgCl_2$ | 1.0 mM spermine | 2.0 M $(NH_4)_2SO_4$ | |
| Both Pos. | 14X | 12 | 0.05 M Na Cacodylate pH 6.5 | 100 mM $MgCl_2$ | 2.0 mM CoH18N6 | 5% isopropanol | |
| **DAPI** | | | | | | | |
| **Motif** | **Binding Ratio** | **Buffer** | **Buffer Components** | | | | |
| Pos. 1 | 3X | 4 | 0.05 M Na Cacodylate pH 6.5 | 18 mM $MgCl_2$ | 2.25 mM spermine | 9% Isopropanol | |
| Pos. 2 (*) | 30X | 20 | 0.05 M HEPES pH 7.5 | 15 mM $MgCl_2$ | 1.0 mM spermidine | 10% dioxane | |
| Both Pos. (*) | 50X | 2 | 0.05 M Cacodylate pH 6.0 | 18 mM $MgCl_2$ | 2.25 mM spermine | 1 mM $CuSO_4$ | 9% Isopropanol |
| **Hoechst** | | | | | | | |
| **Motif** | **Binding Ratio** | **Buffer** | **Buffer Components** | | | | |
| Pos. 1 | 1X | 41 | 0.05 M TRIS pH 8.0 | 200 mM $MgCl_2$ | 15% ethanol | | |
| Pos. 2 | 14X | 40 | 0.05 M Na Cacodylate pH 6.0 | 200 mM $MgCl_2$ | 2.5 M KCl | | |
| Both Pos. (*) | 15X | 40 | 0.05 M Na Cacodylate pH 6.0 | 200 mM $MgCl_2$ | 2.5 M KCl | | |

(*) indicates a temperature ramp rate of -0.2 °C/h was required. All other crystals were obtained using a -0.3 °C/h temperature ramp.

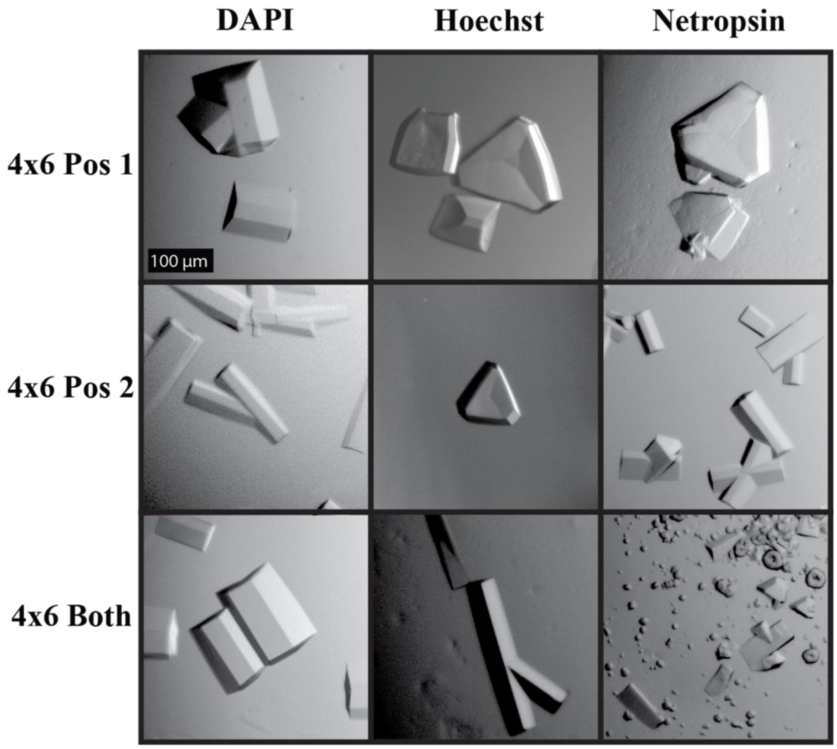

**Supplementary Figure 3. Representative bright field images of the 4x6 MGB co-crystals.** The buffers that correspond to each image are indicated in Supplementary Table 4.

**Supplementary Table 4. Buffer conditions used for the corresponding bright field images**

| 4x6 Motif | | | | | | | |
|---|---|---|---|---|---|---|---|
| **Netropsin** | | | | | | | |
| **Motif** | **Binding Ratio** | **Buffer** | **Buffer Components** | | | | |
| Pos. 1 | 4X | 25 | 0.05 M Na Cacodylate pH 6.5 | 30 mM $MgCl_2$ | 1.0 mM spermine | 1.3 M $Li_2SO_4$ | |
| Pos. 2 | 10X | 28 | 0.05 M Na Cacodylate pH 6.0 | 10 mM $MgCl_2$ | 2.5 mM spermidine | 2.5 M NaCl | |
| Both Pos. | 4X | 39 | 0.05 M TRIS pH 7.5 | 200 mM MgCl2 | 2.5 M NaCl | | |
| **DAPI** | | | | | | | |
| **Motif** | **Binding Ratio** | **Buffer** | **Buffer Components** | | | | |
| Pos. 1 | 20X | 47 | 0.05 M Na cacodylate pH 6.5 | 1.0 mM spermine | 2.0 mM $CoH_{18}N_6$ | 30 mM $CaCl_2$ | 2.0 M LiCl |
| Pos. 2 | 20X | 40 | 0.05 M Na Cacodylate pH 6.0 | 200 mM $MgCl_2$ | 2.5 M KCl | | |
| **Hoechst** | | | | | | | |
| **Motif** | **Binding Ratio** | **Buffer** | **Buffer Components** | | | | |
| Pos. 2 (*) | 1.5X | 40 | 0.05 M Na Cacodylate pH 6.0 | 200 mM $MgCl_2$ | 2.5 M KCl | | |
| Both Pos. | 4X | 13 | 0.05 M TRIS pH 8.0 | 10 mM $MgCl_2$ | 1.0 mM $CoH_{18}N_6$ | 20% ethanol | |

(*) indicates a temperature ramp rate of -0.2 °C/h was required. All other crystals were obtained using a -0.3 °C/h temperature ramp.

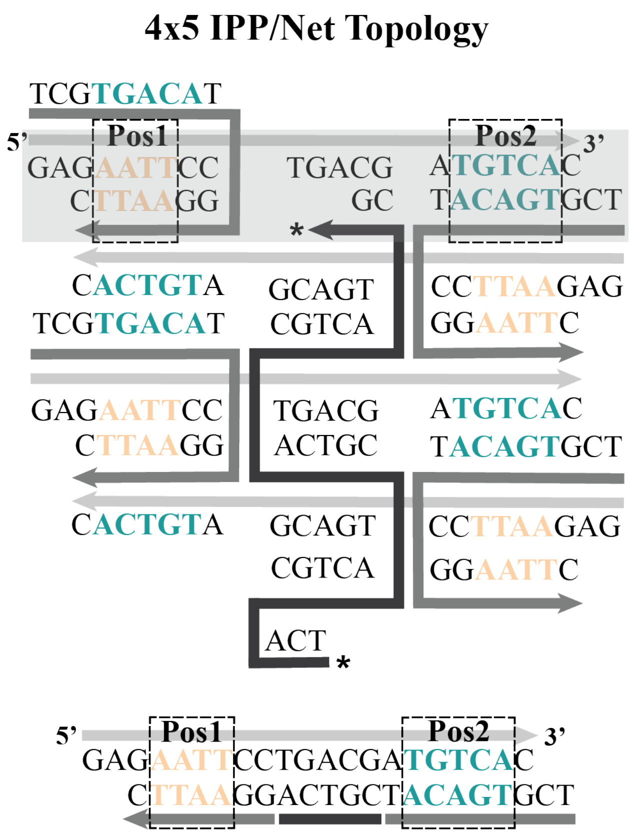

**Supplementary Figure 4. Topological schematic of the 4x5 motif for binding of netropsin and IPP at unique positions.** The netropsin binding sequence (AATT) at Pos1 (tan) and the IPP sequence (TGTCA) at Pos2 (teal) are highlighted. All other features remain the same as the description in Supplementary Figure 1.

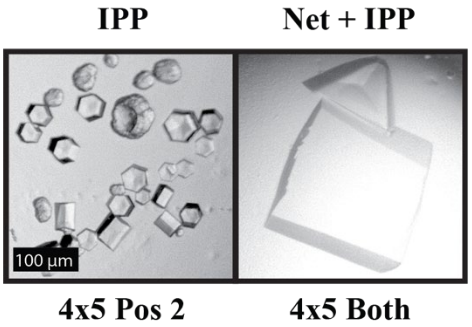

**Supplementary Figure 5. Representative bright field images of the 4x5 IPP Pos2 and IPP Pos2 + netropsin Pos1 co-crystals.**

**Supplementary Table 5. Crystallization buffers corresponding to the bright field images.**

| IPP | | | | | | | | |
|---|---|---|---|---|---|---|---|---|
| **Motif** | **Binding Ratio** | **Buffer** | **Buffer Components** | | | | | |
| Pos. 2 | 7X | 14 | 0.05 M HEPES pH 7.5 | 20 mM $MgCl_2$ | 1.0 mM spermine | 5% PEG 8000 | | |
| IPP + Netropsin | 7X + 2X | 17 | 0.05 M Na cacodylate pH 7.0 | 9 mM $MgCl_2$ | 2.25 mM spermine | 1.8 mM $CoH_{18}N_6$ | 0.9 mM spermidine | 5% PEG 400 |

**Supplementary Table 6. Data collection and refinement statistics for DAPI containing crystals.**

| Variation | DAPI | | | | |
|---|---|---|---|---|---|
| | 4x5 | | | 4x6 | |
| | Pos. 1 | Pos. 2 | Both Pos. | Pos. 1 | Pos. 2 |
| PDB Code | 8TB3 | 8TB4 | 8T7X | 8TA9 | 8TDT |
| **Data Collection** | | | | | |
| Beamline | APS | APS | ALS | APS | APS |
| Space group | $P3_221$ | $P3_2$ | $P3_2$ | $P3_2$ | $P3_2$ |
| Resolution (Å) | 50-3.0 | 50-3.10 | 50-3.05 | 50-2.80 | 50-3.00 |
| **Cell dimensions** | | | | | |
| a, b, c (Å) | 68.6, 68.6, 62.6 | 69.5, 69.5, 59.6 | 68.5, 68.5, 60.1 | 68.4, 68.4, 58.1 | 68.5, 68.5, 56.8 |
| α, β, γ (°) | 90, 90, 120 | 90, 90, 120 | 90, 90, 120 | 90, 90, 120 | 90, 90, 120 |
| Total observations | 25585 | 55936 | 58825 | 70565 | 46135 |
| No. unique reflections | 3620 | 5774 | 5900 | 7272 | 5644 |
| $R_{pim}$ | 0.02 (0.36) | 0.04 (0.63) | 0.03 (0.30) | 0.04 (0.38) | 0.06 (0.83) |
| $CC_{1/2}$ | 1.004 (0.907) | 1.006 (0.644) | 0.912 (0.946) | 1.028 (0.730) | 0.973 (0.806) |
| $I/\sigma I$ | 47.50 (3.75) | 39.00 (0.44) | 49.83 (1.81) | 41.17 (1.21) | 27.28 (1.26) |
| Completeness (%) | 99.8 (100.0) | 99.7 (97.3) | 99.9 (100.0) | 97.2 (76.9) | 95.2 (72.0) |
| Redundancy | 7.1 (7.2) | 9.7 (6.7) | 10.0 (9.2) | 9.7 (7.8)) | 8.2 (6.3) |
| **Refinement** | | | | | |
| $R_{work}/R_{free}$ | 0.1934/0.2397 | 0.1950/0.2379 | 0.1970/0.2218 | 0.1924/0.2440 | 0.2089/0.2257 |
| **No. atoms** | | | | | |
| DNA | 855 | 855 | 855 | 855 | 855 |
| ligand/ion | 0 | 3 | 1 | 2 | 6 |
| solvent | 2 | 0 | 0 | 0 | 4 |
| **R.M.S deviations** | | | | | |
| Bond lengths (Å) | 0.006 | 0.006 | 0.009 | 0.007 | 0.007 |
| Bond angles (°) | 2.002 | 2.066 | 1.088 | 2.329 | 0.891 |

*The value for the highest resolution shell is shown in parentheses

### Supplementary Table 7. Data collection and refinement statistics for Hoechst containing crystals.

| Variation | Hoechst | | | | |
|---|---|---|---|---|---|
| | 4x5 | | | 4x6 | |
| | Pos. 1 | Pos. 2 | Both Pos. | Pos. 2 | Both Pos. |
| PDB Code | 8TB8 | 8TBD | 8T80 | 8TAQ | 8TAM |
| **Data Collection** | | | | | |
| Beamline | ALS | ALS | APS | ALS | ALS |
| Space group | $P3_221$ | $P3_2$ | $P3_2$ | $P3_2$ | R3 |
| Resolution (Å) | 50-2.95 | 50-3.00 | 50-2.90 | 50-2.95 | 50-2.95 |
| **Cell dimensions** | | | | | |
| a, b, c (Å) | 68.2, 68.2, 60.7 | 68.4, 68.4, 58.4 | 68.5, 68.5, 59.1 | 68.9, 68.9, 56.7 | 116.5, 116.5, 49.3 |
| α, β, γ (°) | 90, 90, 120 | 90, 90, 120 | 90, 90, 120 | 90, 90, 120 | 90, 90, 120 |
| Total observations | 67168 | 53271 | 52401 | 63355 | 52906 |
| No. unique reflections | 3640 | 5544 | 6418 | 5913 | 5263 |
| $R_{pim}$ | 0.02 (0.33) | 0.04 (0.41) | 0.05 (0.55) | 0.02 (0.24) | 0.02 (0.40) |
| $CC_{1/2}$ | 0.997 (0.862) | 0.899 (0.742) | 0.981 (0.687) | 1.012 (0.918) | 1.060 (0.822) |
| $I/\sigma I$ | 56.05 (1.60) | 42.89 (0.95) | 37.04 (0.83) | 50.16 (2.33) | 50.44 (1.17) |
| Completeness (%) | 100.0 (100.0 ) | 91.1 (59.8) | 94.4 (66.4) | 93.1 (69.8) | 100.0 (99.3) |
| Redundancy | 18.5 (17.3) | 9.6 (8.1) | 8.2 (5.7) | 10.7 (9.4) | 10.1 (8.7) |
| **Refinement** | | | | | |
| $R_{work}/R_{free}$ | 0.2369/0.2718 | 0.2448/0.2709 | 0.2194/0.2355 | 0.2476/0.2772 | 0.2212/0.2297 |
| **No. atoms** | | | | | |
| DNA | 855 | 853 | 855 | 855 | 855 |
| ligand/ion | 1 | 5 | 1 | 2 | 2 |
| solvent | 2 | 3 | 6 | 3 | 4 |
| **R.M.S deviations** | | | | | |
| Bond lengths (Å) | 0.004 | 0.005 | 0.006 | 0.005 | 0.005 |
| Bond angles (°) | 0.791 | 0.847 | 1.829 | 0.859 | 0.925 |

*The value for the highest resolution shell is shown in parentheses

### Supplementary Table 8. Data collection and refinement statistics for netropsin containing crystals

| Variation | Netropsin | | | | | |
|---|---|---|---|---|---|---|
| | 4x5 | | | 4x6 | | |
| | Pos. 1 | Pos. 2 | Both Pos. | Pos. 1 | Pos. 2 | Both Pos. |
| PDB Code | 8TBO | 8T7B | 8TC2 | 8TAJ | 8TAP | 8TA8 |
| **Data Collection** | | | | | | |
| Beamline | BNL | ALS | ALS | APS | ALS | ALS |
| Space group | $P3_221$ | $P3_2$ | $P3_2$ | $P3_2$ | $P3_2$ | $P3_2$ |
| Resolution (Å) | 50-2.60 | 50-3.05 | 50-3.05 | 50-2.90 | 50-3.10 | 50-3.05 |
| **Cell dimensions** | | | | | | |
| a, b, c (Å) | 68.2, 68.2, 63.1 | 68.8, 68.8, 59.9 | 68.5, 68.5, 59.7 | 68.2, 68.2, 57.9 | 68.6, 68.6, 56.5 | 68.5, 68.5, 58.3 |
| α, β, γ (°) | 90, 90, 120 | 90, 90, 120 | 90, 90, 120 | 90, 90, 120 | 90, 90, 120 | 90, 90, 120 |
| Total observations | 91099 | 59853 | 59614 | 61699 | 44258 | 53988 |
| No. unique reflections | 5449 | 5937 | 5931 | 6234 | 4797 | 5425 |
| $R_{pim}$ | 0.02 (0.35) | 0.020 (0.208) | 0.03 (0.27) | 0.037 (0.341) | 0.03 (0.41) | 0.02 (0.26) |
| $CC_{1/2}$ | 0.956 (0.755) | 0.982 (0.910) | 0.954 (0.854) | 0.976 (0.868) | 0.982 (0.751) | 0.990 (0.922) |
| $I/\sigma I$ | 66.14 (1.64) | 48.81 (1.75) | 42.13 (1.47) | 37.59 (1.29) | 31.04 (0.682) | 48.38 (1.53) |
| Completeness (%) | 99.1 (96.9) | 98.5 (83.8) | 99.8 (96.3) | 9.9 (8.5) | 89.4 (52.0) | 93.2 (74.4) |
| Redundancy | 16.7 (12.0) | 10.1 (8.2) | 10.1 (8.6) | 93.3 (70.4) | 9.2 (5.8) | 8.3 (10.0) |
| **Refinement** | | | | | | |
| $R_{work}/R_{free}$ | 0.2119/0.2430 | 0.1917/0.2108 | 0.2150/0.2605 | 0.2141/0.2243 | 0.2069/0.2517 | 0.2146/0.2426 |
| **No. atoms** | | | | | | |
| DNA | 855 | 855 | 855 | 855 | 855 | 855 |
| ligand/ion | 4 | 0 | 3 | 2 | 0 | 0 |
| solvent | 3 | 0 | 8 | 0 | 2 | 2 |
| **R.M.S deviations** | | | | | | |
| Bond lengths (Å) | 0.010 | 0.011 | 0.007 | 0.006 | 0.006 | 0.006 |
| Bond angles (°) | 1.120 | 1.959 | 0.837 | 0.854 | 0.920 | 0.860 |

*The value for the highest resolution shell is shown in parentheses

**Supplementary Table 9. Data collection and refinement statistics for IPP and IPP + netropsin crystals**

| Variation | ImPyPy | ImPyPy + Netropsin |
|---|---|---|
| | 4x5 | |
| | Pos. 2 | Both Pos. |
| PDB Code | 8TC4 | 8TC6 |
| **Data Collection** | | |
| Beamline | ALS | BNL |
| Space group | $P3_2$ | $P3_221$ |
| Resolution (Å) | 50-3.05 | 50-2.45 |
| **Cell dimensions** | | |
| a, b, c (Å) | 68.7, 68.7, 62.2 | 67.9, 67.9, 65.5 |
| α, β, γ (°) | 90, 90, 120 | 90, 90, 120 |
| Total observations | 50178 | 125179 |
| No. unique reflections | 5432 | 6739 |
| $R_{pim}$ | 0.02 (0.58) | 0.02 (0.35) |
| $CC_{1/2}$ | 1.008 (0.628) | 1.114 (0.781) |
| $I/\sigma I$ | 35.69 (0.64) | 64.00 (2.00) |
| Completeness (%) | 87.2 (54.2) | 100.0 (100.0) |
| Redundancy | 9.2 (8.3) | 18.6 (16.6) |
| **Refinement** | | |
| $R_{work}/R_{free}$ | 0.2289/0.2535 | 0.2721/0.2942 |
| **No. atoms** | | |
| DNA | 855 | 855 |
| ligand/ion | 2 | 2 |
| solvent | 1 | 4 |
| **R.M.S deviations** | | |
| Bond lengths (Å) | 0.006 | 0.006 |
| Bond angles (°) | 0.813 | 0.834 |

*The value for the highest resolution shell is shown in parentheses

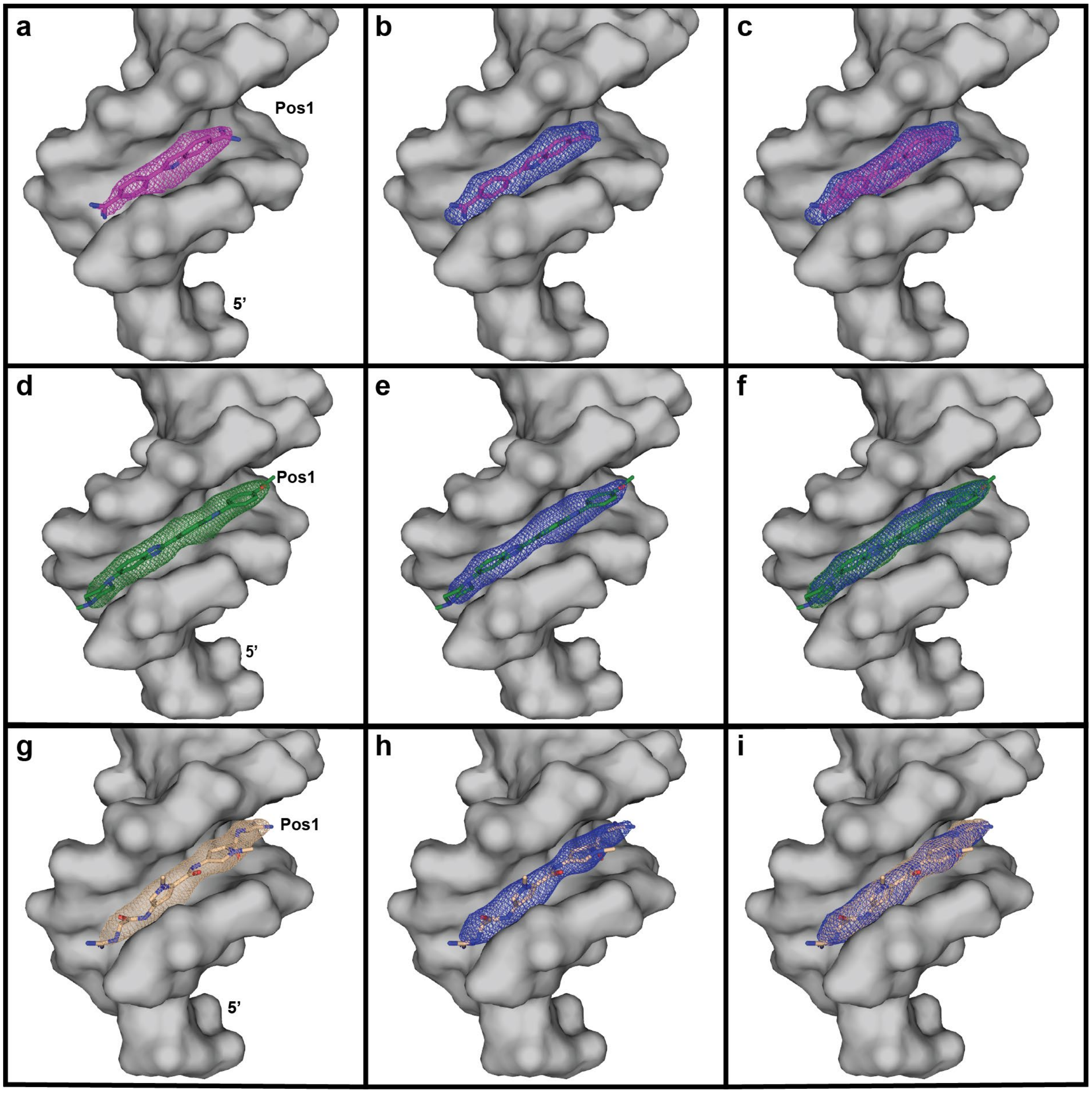

**Supplementary Figure 6. Electron density maps and structures of MGBs bound at Pos1 in the 4x5 motif.** (a) DAPI (purple) $2F_o - F_c$ map (purple) contoured at $\sigma = 1.2$ (b) DAPI polder map (blue) contoured at $\sigma = 3.0$ (c) Superposition of the $2F_o - F_c$ and polder maps in (a) and (b); (d) Hoechst (green) $2F_o - F_c$ map (green) contoured at $\sigma = 1.2$ (e) Hoechst polder map (blue) contoured at $\sigma = 3.0$ (f) Superposition of the $2F_o - F_c$ and polder maps in (d) and (e); (g) netropsin (tan) $2F_o - F_c$ map (tan) contoured at $\sigma = 1.2$ (h) netropsin polder map (blue) contoured at $\sigma = 3.0$ (i) Superposition of the $2F_o - F_c$ and polder maps in (g) and (h).

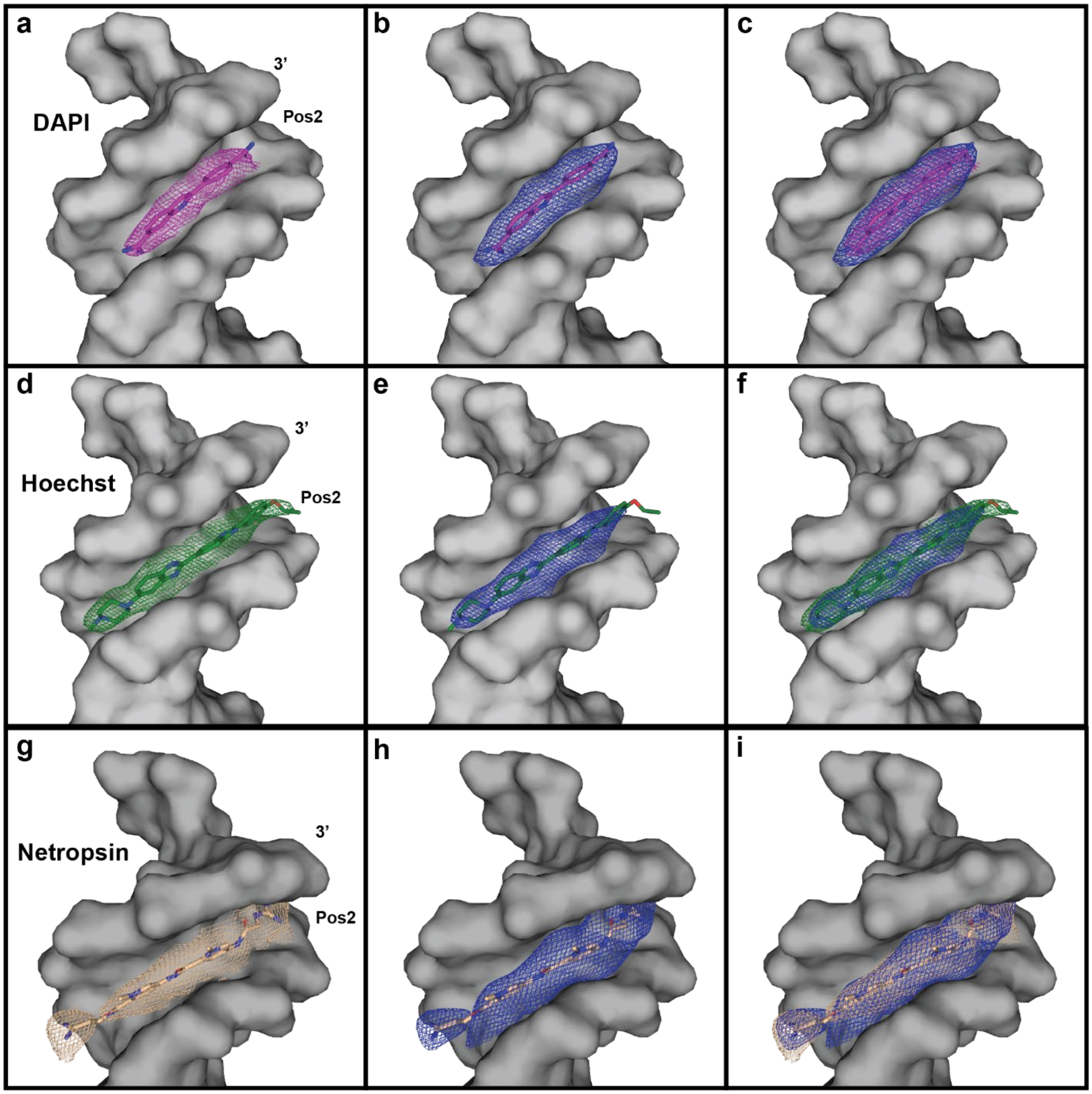

**Supplementary Figure 7. Electron density maps and structures of MGBs bound at Pos2 in the 4x5 motif.** (a) DAPI (purple) $2F_o - F_c$ map (purple) contoured at $\sigma = 1.2$ (b) DAPI polder map (blue) contoured at $\sigma = 2.8$ (c) Superposition of the $2F_o - F_c$ and polder maps in (a) and (b); (d) Hoechst (green) $2F_o - F_c$ map (green) contoured at $\sigma = 1.2$ (e) Hoechst polder map (blue) contoured at $\sigma = 3.0$ (f) Superposition of the $2F_o - F_c$ and polder maps in (d) and (e); (g) netropsin (tan) $2F_o - F_c$ map (tan) contoured at $\sigma = 0.8$ (h) netropsin polder map (blue) contoured at $\sigma = 2.6$ (i) Superposition of the $2F_o - F_c$ and polder maps in (g) and (h).

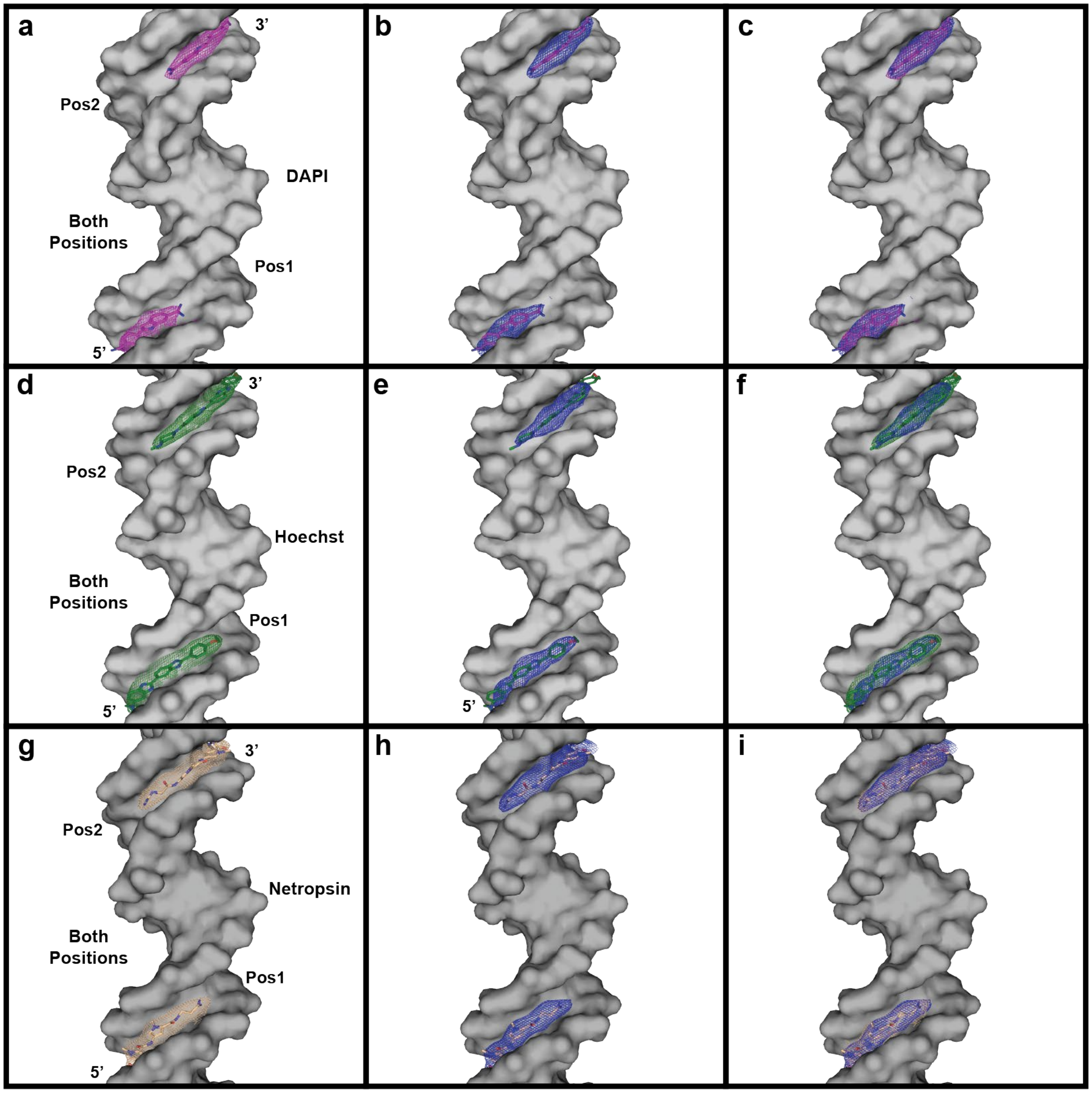

**Supplementary Figure 8. Electron density maps and structures of MGBs bound at Pos2 in the 4x5 motif.** (a) DAPI (purple) Pos1 and Pos2 $2F_o - F_c$ maps (purple) contoured at $\sigma = 1.2$; (b) DAPI polder map (blue) contoured at $\sigma = 2.8$ for both positions; (c) superposition of the $2F_o - F_c$ and polder maps in (a) and (b); (d) Hoechst (green) Pos1 and Pos2 $2F_o - F_c$ maps (green) contoured at $\sigma = 1.0$; (e) Hoechst polder map (blue) contoured at $\sigma = 3.0$ for both positions; (f) superposition of the $2F_o - F_c$ and polder maps in (d) and (e); (g) netropsin (tan) Pos1 and Pos2 $2F_o - F_c$ maps (tan) contoured at $\sigma = 1.6$; (h) netropsin polder map (blue) contoured at $\sigma = 2.8$ for both positions; and (i) superposition of the $2F_o - F_c$ and polder maps in (g) and (h).

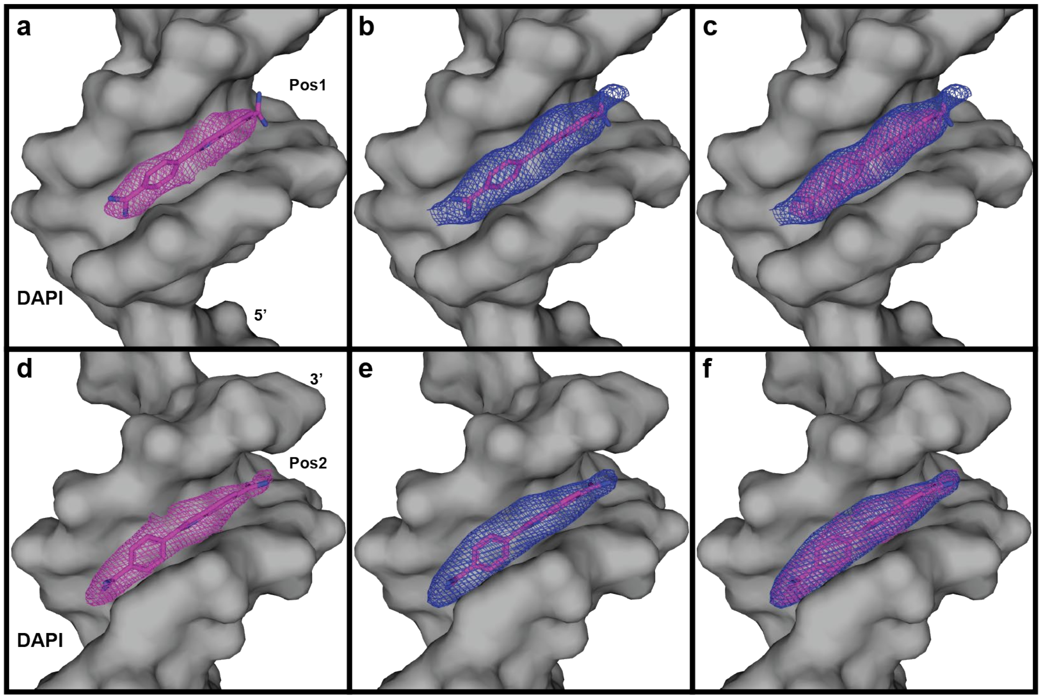

**Supplementary Figure 9. Electron density maps and structures of DAPI at Pos1 and Pos2 in the 4x6 motif.** (a) Pos1 DAPI (purple) $2F_o - F_c$ map (purple) contoured at $\sigma = 1.1$ (b) DAPI polder map (blue) contoured at $\sigma = 2.8$; (c) Superposition of the $2F_o - F_c$ and polder maps in (a) and (b); (d) Pos2 DAPI (purple) $2F_o - F_c$ map (purple) contoured at $\sigma = 1.4$; (b) DAPI polder map (blue) contoured at $\sigma = 3.5$; and (c) Superposition of the $2F_o - F_c$ and polder maps in (d) and (e). Note: Adequate electron density was not achieved to accurately solve the BP DAPI structure in the 4x6 system.

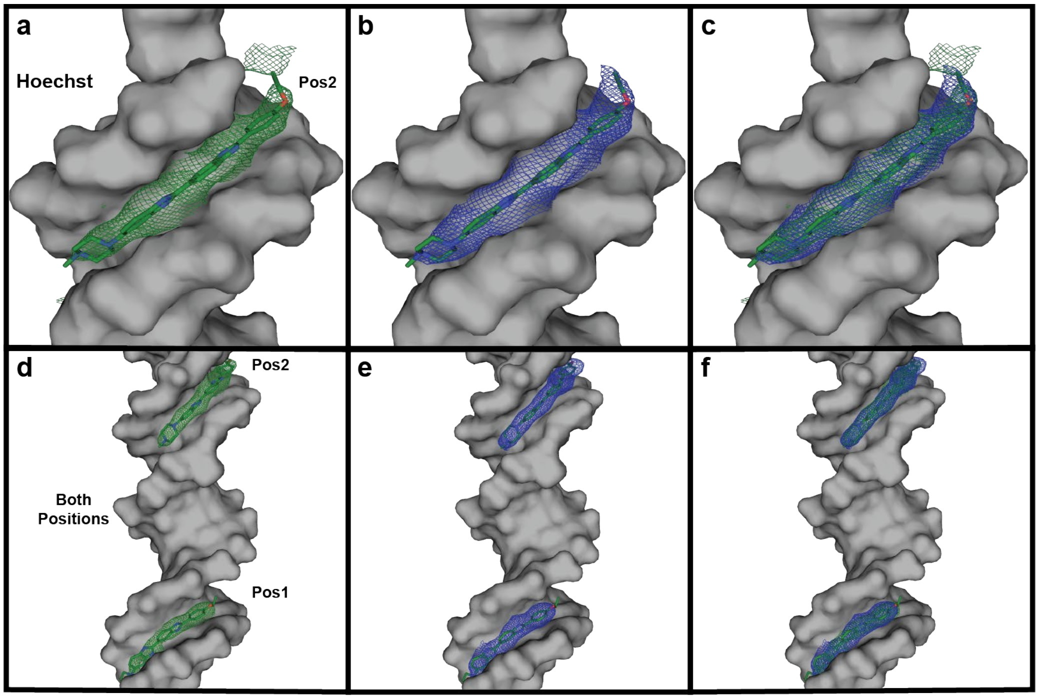

**Supplementary Figure 10. Electron density maps and structures of Hoechst molecules bound at Pos2 and BP in the 4x6 motif.** (a) Pos2 Hoechst (green) $2F_o - F_c$ map (green) contoured at $\sigma = 1.2$; (b) DAPI polder map (blue) contoured at $\sigma = 2.4$; and (c) Superposition of the $2F_o - F_c$ and polder maps in (a) and (b). The additional peak in the map (a) on the ethoxyphenyl end of the molecule is likely attributable to noise inherent to the map. The polder map (b) substantiates that the molecule was built properly; (d) Hoechst (green) Pos1 and Pos2 $2F_o - F_c$ maps (green) contoured at $\sigma = 1.3$; (e) Hoechst polder map (blue) contoured at $\sigma = 3.0$ for both positions; and (f) superposition of the $2F_o - F_c$ and polder maps in (d) and (e). Note: Adequate electron density was not achieved to accurately solve the Pos1 Hoechst structure in the 4x6 system.

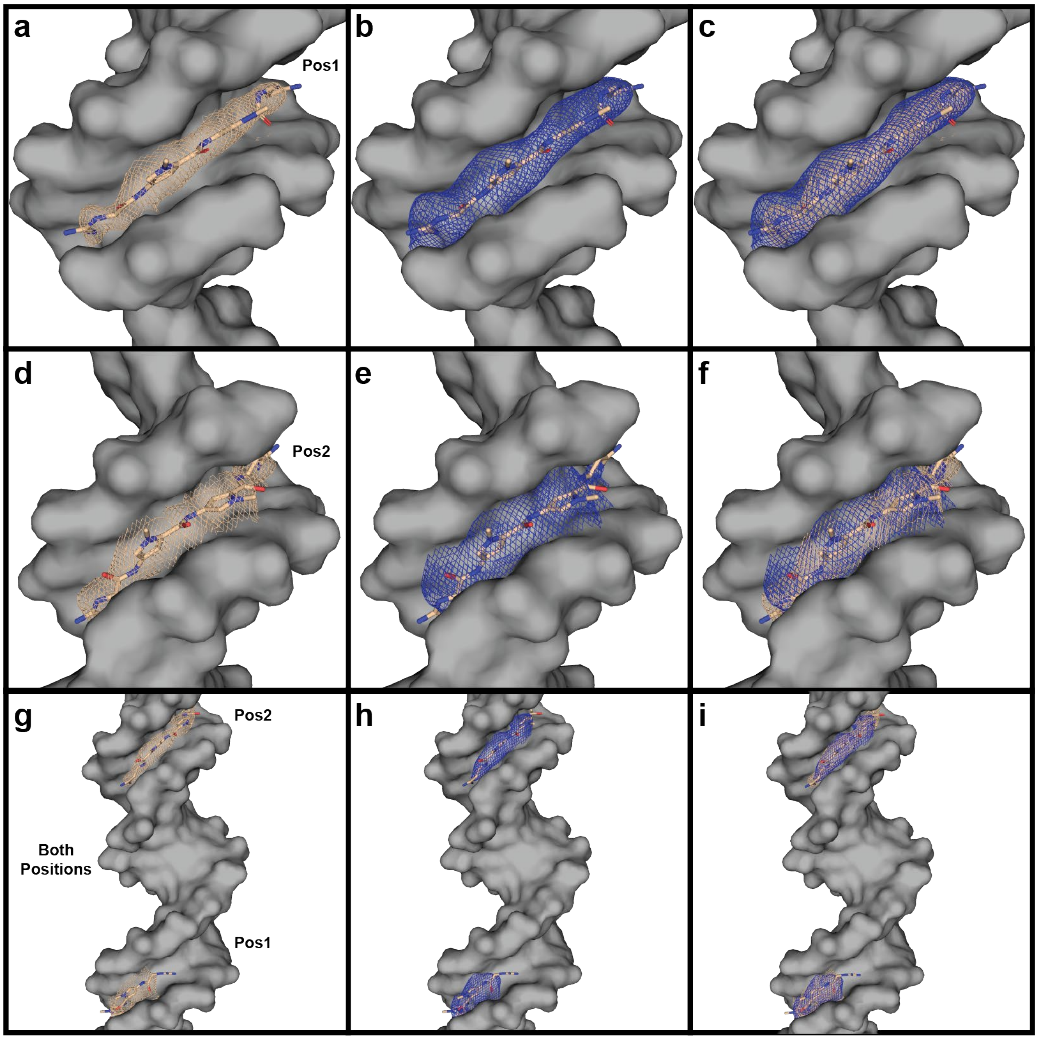

**Supplementary Figure 11. Electron density maps and structures of netropsin bound at Pos1, Pos2 and BP in the 4x6 motif.** (a) Pos1 netropsin (tan) $2F_o - F_c$ map (tan) contoured at $\sigma = 1.2$; (b) netropsin polder map (blue) contoured at $\sigma = 2.6$; (i) Superposition of the $2F_o - F_c$ and polder maps in (a) and (b); (d) Pos2 netropsin (tan) $2F_o - F_c$ map (tan) contoured at $\sigma = 0.8$ (e) netropsin polder map (blue) contoured at $\sigma = 2.6$ (f) Superposition of the $2F_o - F_c$ and polder maps in (d) and (e). (g) netropsin (tan) Pos1 $2F_o - F_c$ map (tan) contoured at $\sigma = 1.4$ and Pos2 $2F_o - F_c$ map at $\sigma = 0.8$; (h) netropsin polder map (blue) contoured at $\sigma = 2.5$ for both positions; and (i) superposition of the $2F_o - F_c$ and polder maps in (g) and (h).

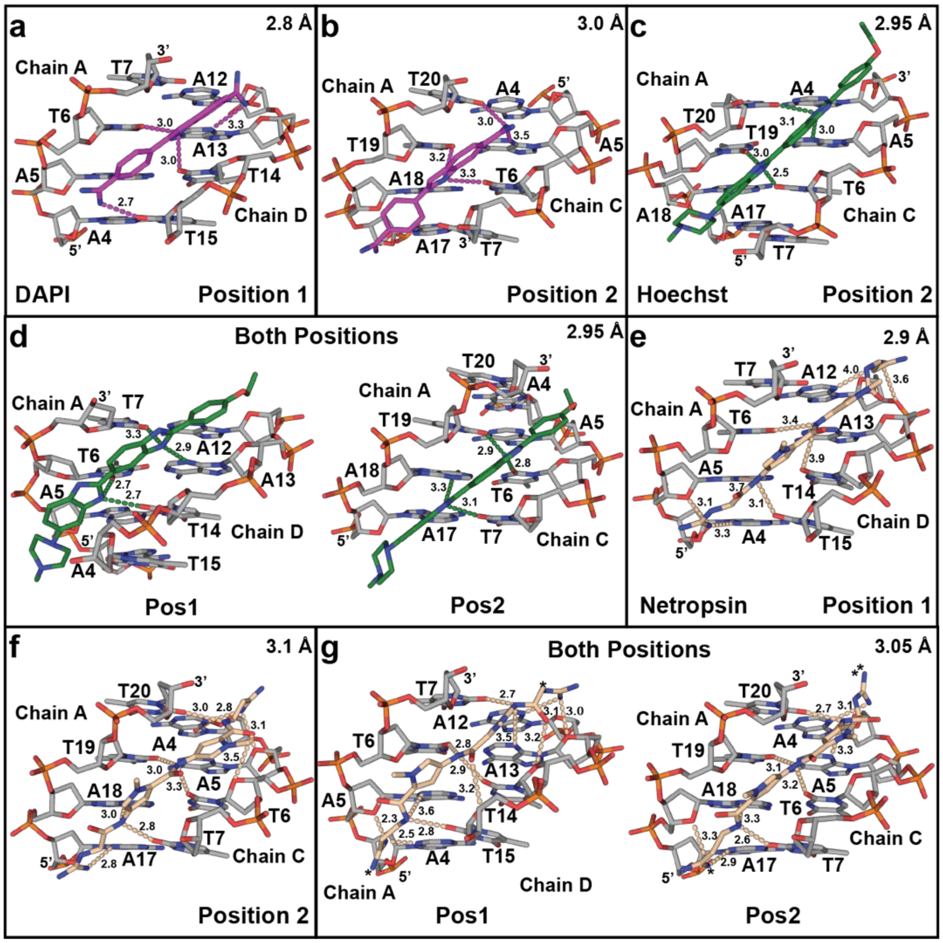

**Supplementary Figure 12. Structure and electrostatic coordination of DAPI, Hoechst, and netropsin to the AATT minor groove in the 4x6 lattices.** The AATT sites of each minor groove are shown in the 4x6 lattice. The chains containing each binding sequence are shown for all structures, and the 5' and 3' ends are indicated to orient the direction of the sequence in one of the component chains with the nucleotide identity and number within each chain shown. Atoms are colored as follows: DNA carbons (gray), oxygen (red), nitrogen (blue), phosphate (orange). MGB carbons: DAPI (purple), Hoechst (green), and netropsin (tan). All electrostatic contacts are indicated with dashed lines colored to match the MGB, and all hydrogen bonding distances between the molecule and the DNA are shown. Distances of strong (2.5 Å) to weak (4.0 Å) electrostatic interactions were used as the criteria for a proper contact and are shown with dashes. (a) Electrostatic coordination scheme of DAPI (purple) Pos1 minor groove; (b) DAPI (purple) Pos2 minor groove; (c) Electrostatic coordination scheme of Hoechst 33342 (green) at Pos2; (d) Hoechst 33342 (green) at both positions; (e) Electrostatic coordination scheme of netropsin (tan) Pos1 minor groove with (f) netropsin (tan) Pos2 minor groove; (g) netropsin (tan) at both positions. The resulting resolutions for each structure are shown in the upper right of each panel. Asterisks indicate additional backbone interactions outside the AATT pocket. Note: Adequate electron density was not achieved to accurately solve the structure of the BP DAPI, and Pos1 Hoechst structures in the 4x6 system.

**Supplementary Table 10. Structure alignment of DAPI, Hoechst, and netropsin (1D30, 129D, and 6BNA)**

| 4x5 MGB structural agreement (RMSD) | | | | |
|---|---|---|---|---|
| MGB | Position 1 | Position 2 | Both Positions | |
| | | | Position 1 | Position 2 |
| Netropsin | 0.934 | 0.628 | 0.379 | 0.398 |
| DAPI | 0.128 | 1.404 | 0.125 | 0.215 |
| Hoechst 33342 | 0.43 | 0.472 | 0.377 | 0.539 |

| 4x6 MGB structural agreement (RMSD) | | | | |
|---|---|---|---|---|
| MGB | Position 1 | Position 2 | Both Positions | |
| | | | Position 1 | Postion 2 |
| Netropsin | 0.146 | 0.408 | 0.962 | 0.351 |
| DAPI | 1.32 | 0.276 | | |
| Hoechst 33342 | | 0.472 | 0.299 | 0.558 |

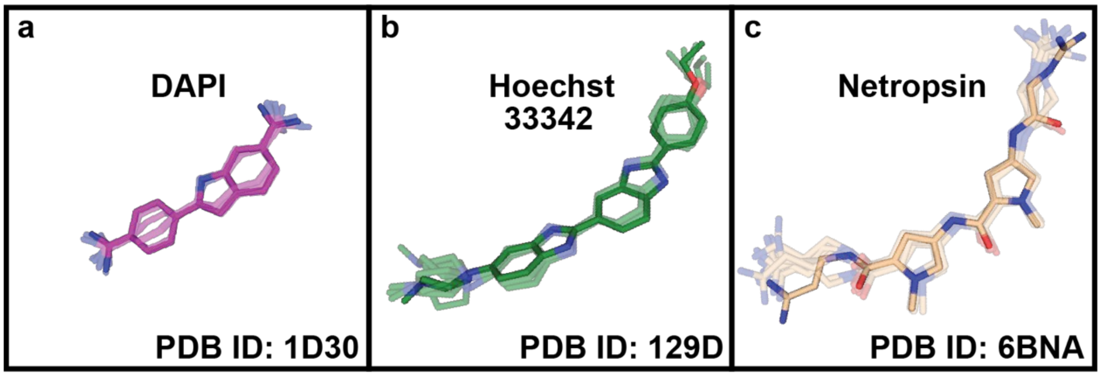

**Supplementary Figure 13. Reference molecule superpositions of MGB structures to original crystal structures.** All crystal structures solved in both the 4x5 and 4x6 systems have been superimposed as translucent stick representations with the respective opaque structures. (a) All DAPI structures (translucent) superimposed with 1D30 (solid); (b) All Hoechst 33342 structures (translucent) superimposed with 129D (solid); and (c) All netropsin structures (translucent) superimposed with 6BNA (solid). The labile propylamidinium and guanidinium moieties on either end of 6BNA show a slight departure from the other structures; however, all minor groove binding amides are in excellent agreement.

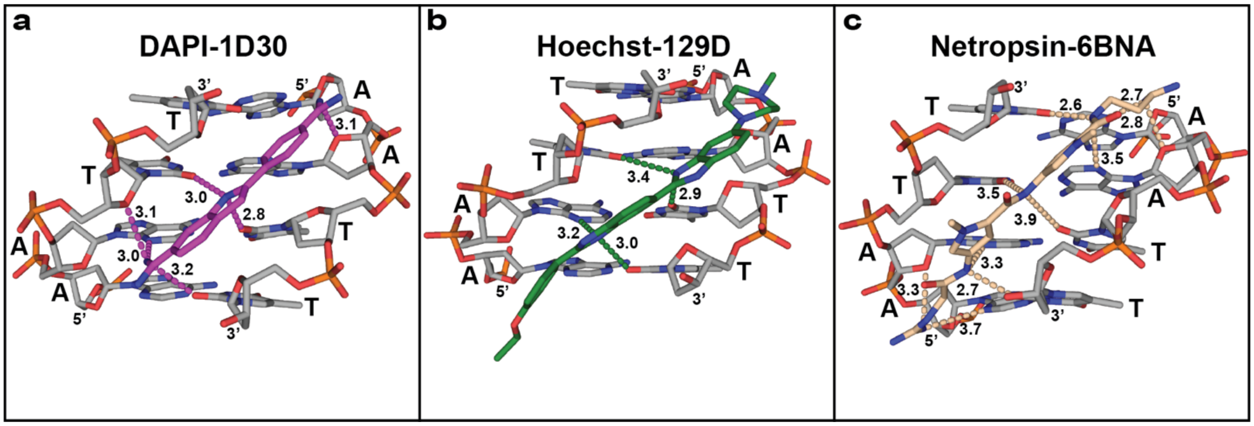

**Supplementary Figure 14. Structure and electrostatic coordination of DAPI (1D30), Hoechst (129D), and netropsin (6BNA) to the AATT minor groove in the original structures.** The AATT sites of each minor groove are shown with bases labeled, and the 5' and 3' ends are indicated. Atoms are colored as follows: DNA carbons (gray), oxygen (red), nitrogen (blue), and phosphate (orange). MGB carbons: DAPI (purple), Hoechst (green), and netropsin (tan). All electrostatic contacts are indicated with dashed lines colored to match the MGB, and all hydrogen bonding distances between the molecule and the DNA are shown. The individual PDB accession codes are labeled for each original structure. (a) DAPI (PDB: 1D30); all hydrogen bonds are indicated including those with neighboring sugar moieties, but only contacts between the molecule and the MG bases are indicated in Supplementary Figure 14 (below); (b) Hoechst (PDB: 129D); and (c) netropsin (PDB: 6BNA).

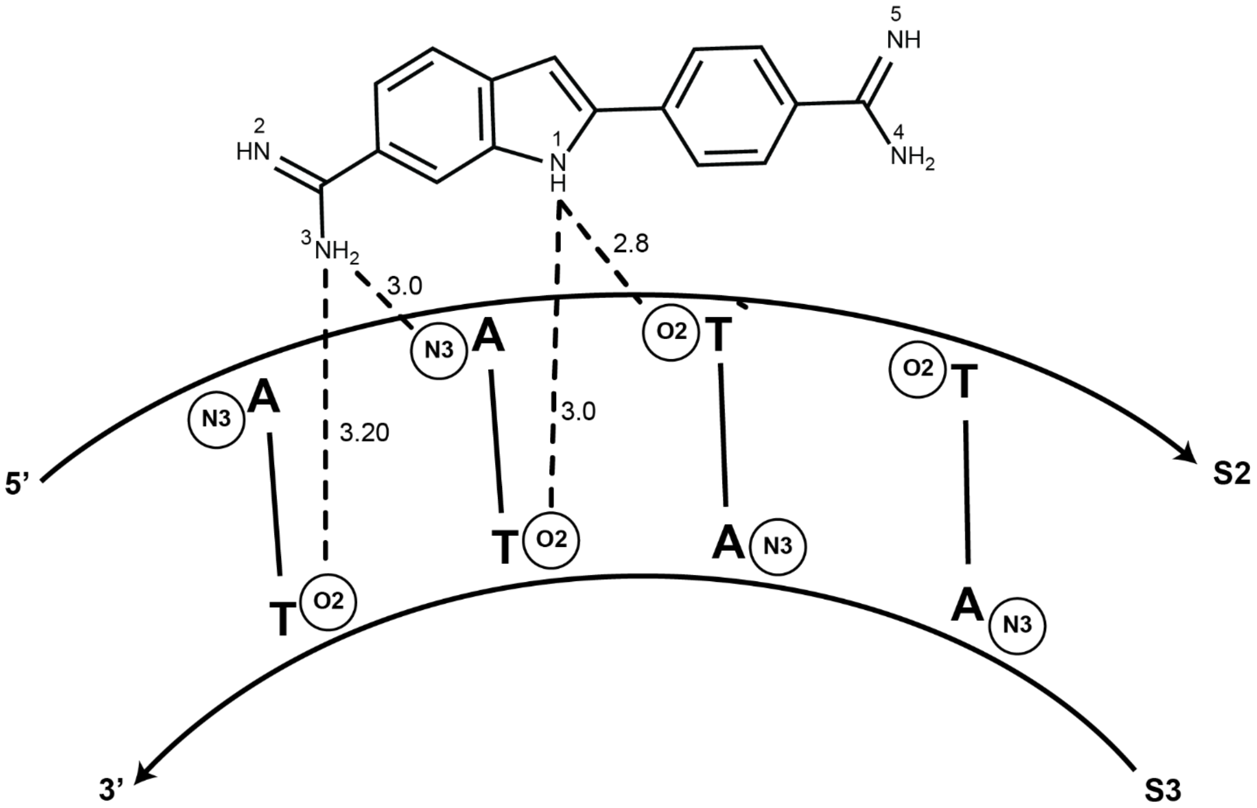

**Supplementary Figure 15. 2D schematic of the DAPI-AATT polar contacts discovered in PDB 1D30.** The original crystal structure revealed four main electrostatic interactions that included a bifurcated hydrogen bond from the indole nitrogen (N1) to two T(O2) atoms immediately diagonal from one another, and from the N3-amine group on the 5' amidine moiety to an A(N3) and the (T)O2 on the 3' end of S3. Each respective hydrogen bonding distance is indicated, and the oligonucleotides are named according to the convention used in this work. The 5' and 3' directionality of the constituent strands is also indicated.

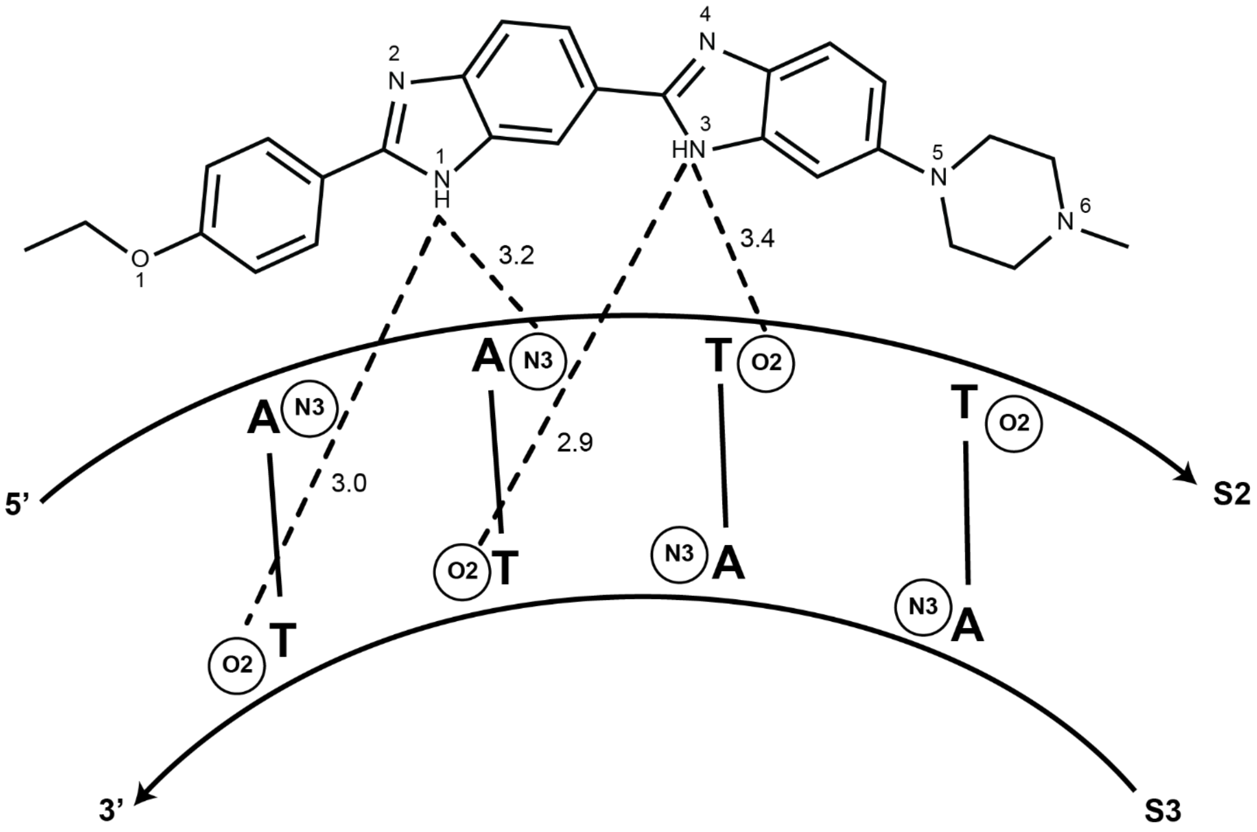

**Supplementary Figure 16. 2D schematic of the Hoechst-AATT polar contacts discovered in PDB 129D.** The original crystal structure revealed four main electrostatic interactions that included a three-centered hydrogen bond from the imidazole N3 to a T(O2) and a T(O2) diagonal to it, and from a bifurcated hydrogen bond from the imidazole N3 on the neighboring benzimidazole to the A(N3) on the opposite side, and to a T(O2) one base downstream from it. Each respective hydrogen bonding distance is indicated, and the oligonucleotides are named according to the convention used in this work. The 5’ and 3’ directionality of the constituent strands is also indicated.

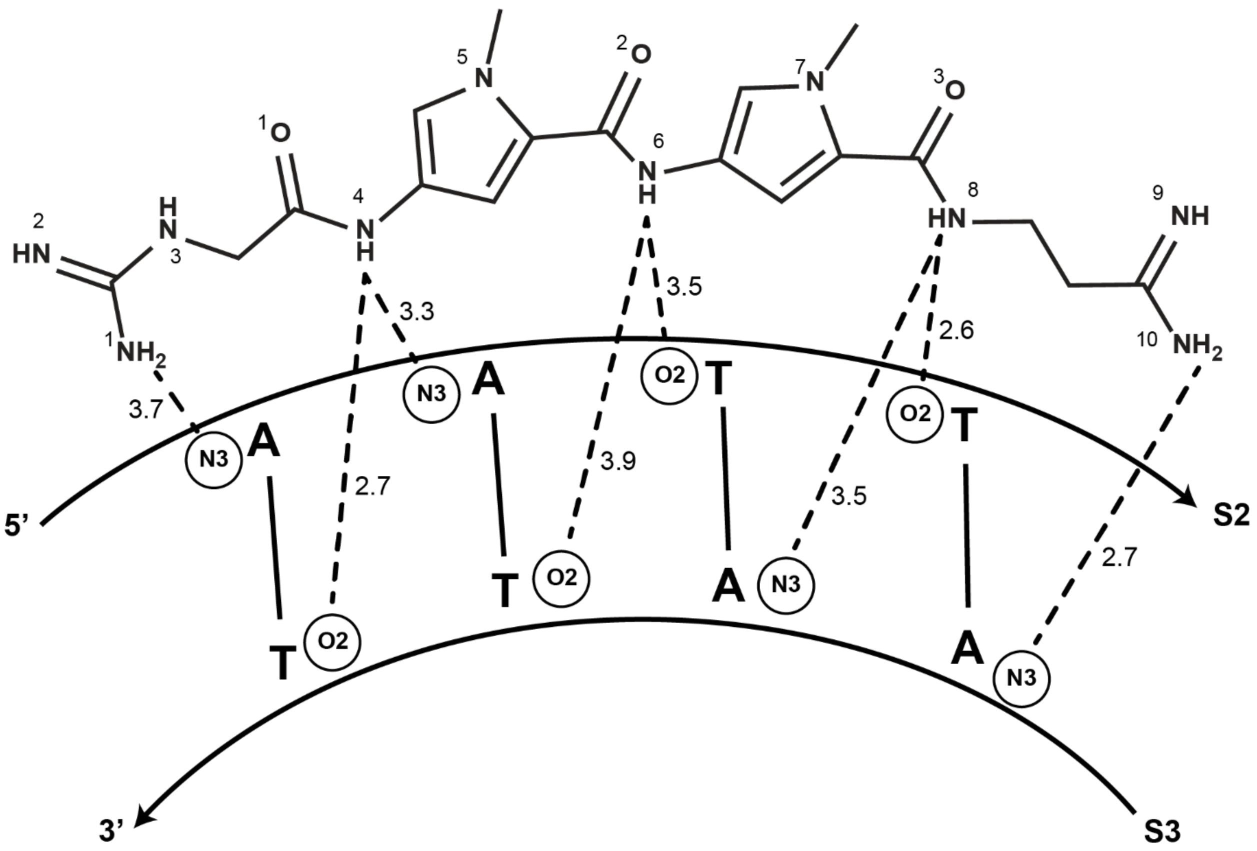

**Supplementary Figure 17. 2D schematic of the netropsin-AATT polar contacts discovered in PDB 6BNA.** The original netropsin structure revealed eight main electrostatic interactions and are indicated with dashed lines. Specifically, the N1 of the 5'-guanidinium group interacts with the 5'-A(N3) and from the N4 amide to its pairing T(O2) and the second 5'-A(N3). The amide connecting the two N-methylated pyrrole rings forms a three centered hydrogen bond to T-O2 groups on opposing sides. The N8 amide moiety interacts with the 5'-T(O2) and an A(N3) on the opposing S3 strand. A final interaction is made between the N10 amine from the propylamidinium group on the 3' end of netropsin. Each respective hydrogen bonding distance is indicated, and the oligonucleotides are named according to the convention used in this work. The 5' and 3' directionality of the constituent strands is also indicated.

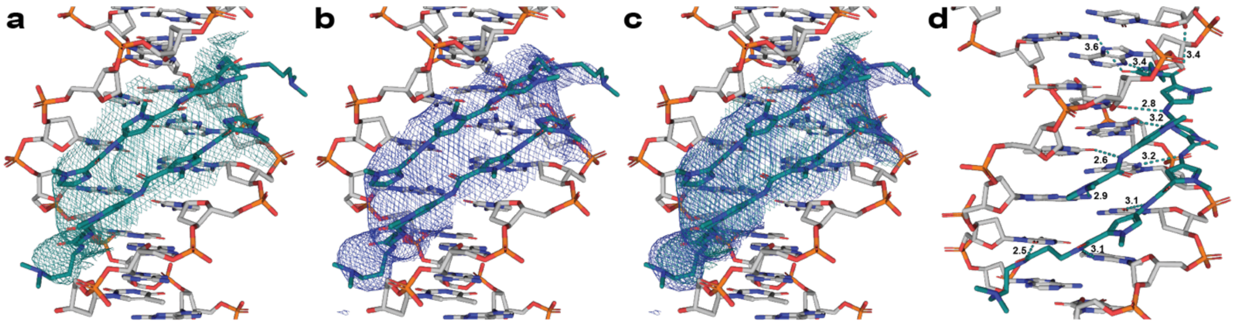

**Supplementary Figure 18. Electron density and polar contacts of the IPP Pos2 structure.** (a) Pos2 IPP (teal) $2F_o - F_c$ map (teal) contoured at $\sigma = 1.0$; (b) DAPI polder map (blue) contoured at $\sigma = 2.4$; and (c) Superposition of the $2F_o - F_c$ and polder maps in (a) and (b). The flexible tail on the top monomer is not accounted for; however, the polder map revealed the detail for proper model building. (d) Mapped electrostatic interactions are indicated with dashed lines between the antiparallel homodimer and the DNA bases. The main interactions and distances are consistent with the structure described in the main text. Atoms are colored as follows: IPP carbons (teal), DNA carbons (gray), oxygen (red), nitrogen (blue), and phosphate (orange).

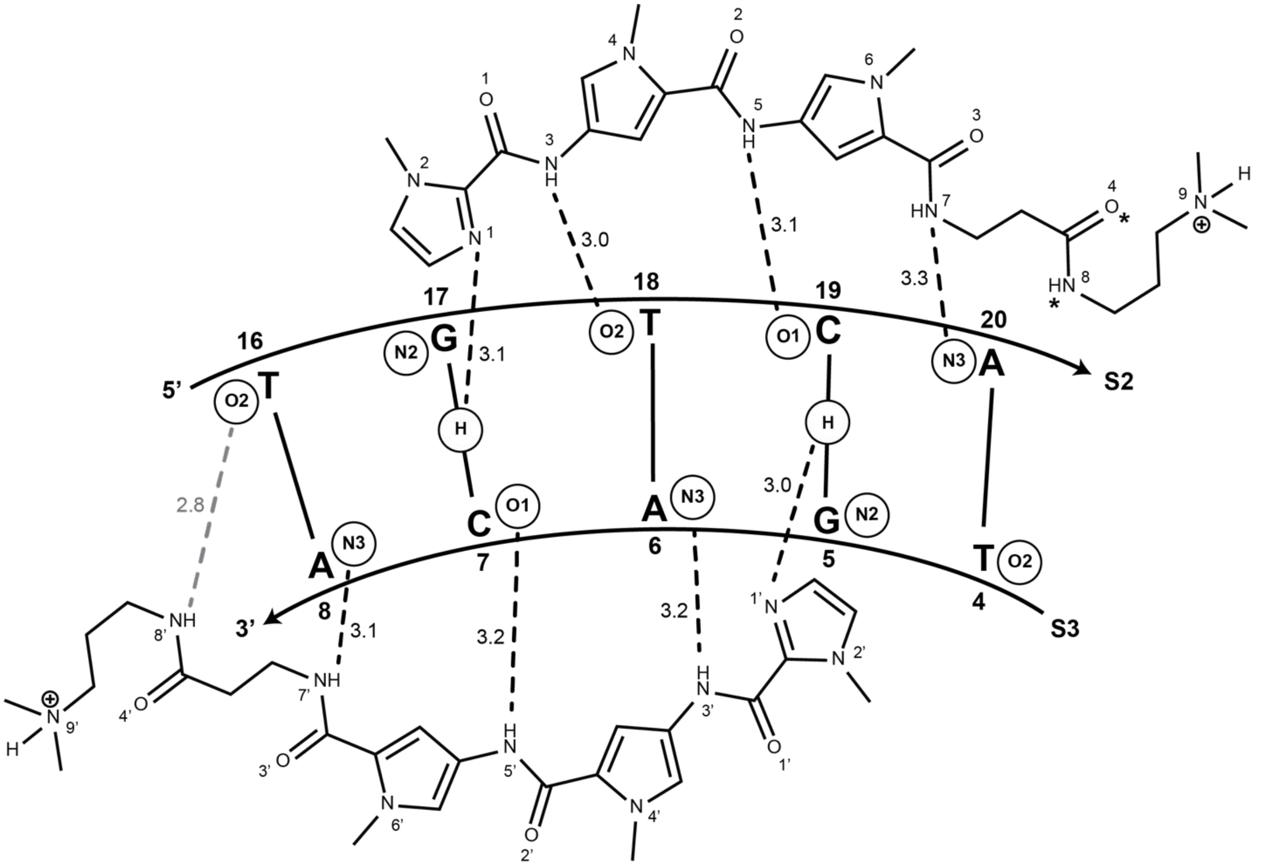

**Supplementary Figure 19. 2D schematic of the IPP-TGTCA polar contacts in the 4x5 IPP + netropsin co-crystal.** Schematic diagram of the polar interactions and distances between the IPP and the minor groove. The 5’ → 3’ directionality of strand S2 carries the TGTCA sequence. The IPP pairs (top to bottom in this direction are as follows Im/Py where N1 hydrogen bonds with N2 hydrogen of G17 and the carboxamide on the carboxyl side of the Py contacts the O1 of C7. The degenerate Py/Py pair forms amide contacts with T18•A6 O2 and N3, respectively. When the Py/Py stacks the amide shows no preference for A•T/T•A, but does not recognize G•C/C•G pairs. The Py/Im pair shows exclusive preference for C•G pairs where the carboxamide (N5) on the C-terminal side of the pyrrole hydrogen bonds with the O1 of C19, and the N1’ of the imidazole contacts the hydrogen of the N2 of G5. The N7 and N7’ hydrogen bond with the N3 of A20 and A8 on the 3’ ends of the binding pocket. The 8’ amide on the N-terminal β-Dp tail forms an additional contact with T16-O2. Asterisks at O4 and N8 indicate two additional tail contacts with were also identified between the N7 amide moiety on the opposing tails of each IPP monomer to adenine(N3) on each 3’ end of the sequence. All hydrogen bonding distances that correspond to the crystal structure are shown.

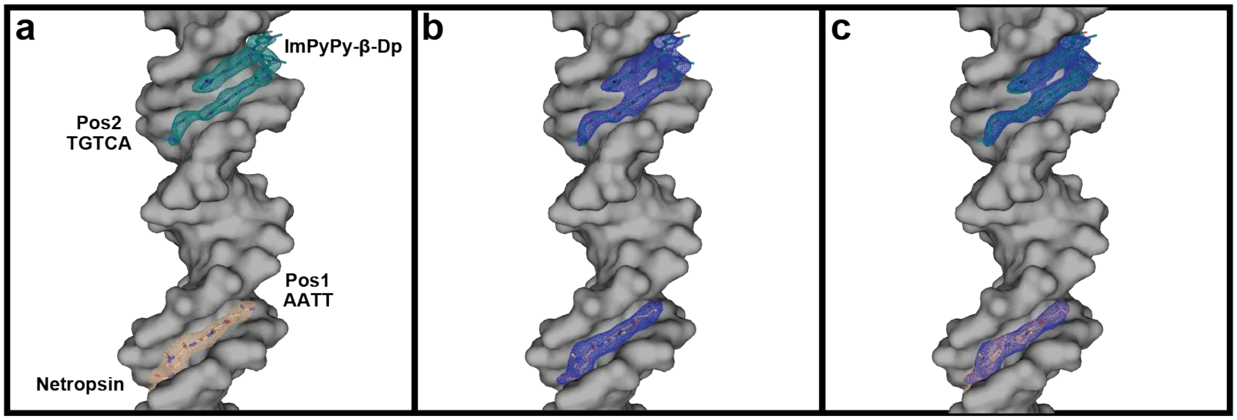

**Supplementary Figure 20. Electron density containing the structures of IPP and netropsin in the 4x5 motif.** (a) Pos2 IPP (teal) $2F_o - F_c$ map (teal) contoured at $\sigma = 1.5$, and netropsin (tan) $2F_o - F_c$ map (tan) contoured at $\sigma = 1.0$. The AATT (Pos1) and TGTCA (Pos2) are labeled at each respective binding site; (b) Pos2 polder map (blue) contoured at $\sigma = 2.4$ and Pos1 polder map (blue) contoured at $\sigma = 2.8$; and (c) Superposition of the $2F_o - F_c$ and polder maps in (a) and (b). Atoms are colored as follows: IPP carbons (teal), oxygen (red), and nitrogen (blue).

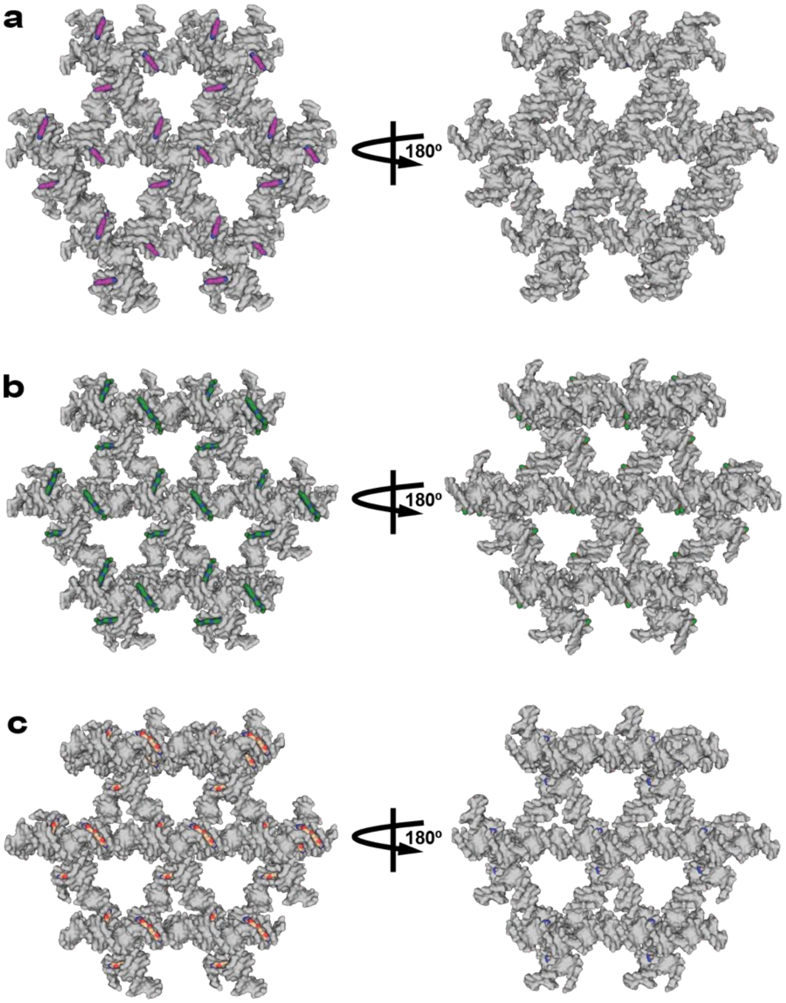

**Supplementary Figure 21. 4x5 Pos2 crystal lattice packing and molecule orientation.** (a) DAPI (purple) Pos2; (b) Hoechst (green) Pos2; (c) netropsin Pos2. Each $P3_2$ lattice is rendered as a surface cross-section of four consecutive layers in the crystal with the AATT minor groove binding sequences at Pos2 in each helical array where each MGB molecule is attached in the structure. The $P3_2$ lattices present all bound MGBs on the front face of the crystal along each constituent duplex in the helical array. When the crystal is oriented 180° no minor grooves on the reverse side contain the bound molecule. The Pos2 lattice results in 50% occupied crystal. The surface of the DAPI (purple carbons), Hoechst (purple carbons), and netropsin (tan carbons) molecules have been surface rendered, with oxygen (red) and nitrogen (blue) also represented.

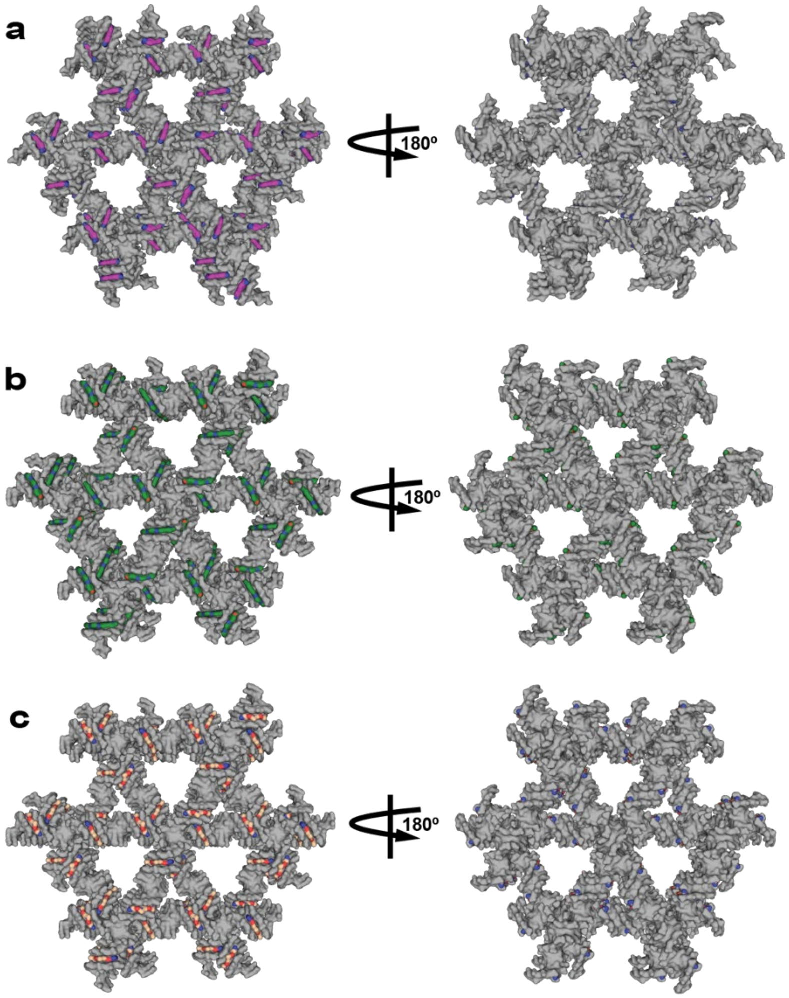

**Supplementary Figure 22. 4x5 crystal lattice packing and orientation at both positions.** (a) DAPI (purple) BP; (b) Hoechst (green) BP; (c) netropsin BP. Each $P3_2$ lattice is rendered as a surface cross-section of four consecutive layers in the crystal with the AATT minor groove binding sequences at BP in each helical array where each MGB molecule is attached in the structure. The $P3_2$ lattices present all bound MGBs on the front face of the crystal along each constituent duplex in the helical array. When the crystal is oriented 180° no minor grooves on the reverse side contain the bound molecule. The BP lattice results in 100% occupied crystal. The surface of the DAPI (purple carbons), Hoechst (purple carbons), and netropsin (tan carbons) molecules have been surface rendered, with oxygen (red) and nitrogen (blue) also represented.

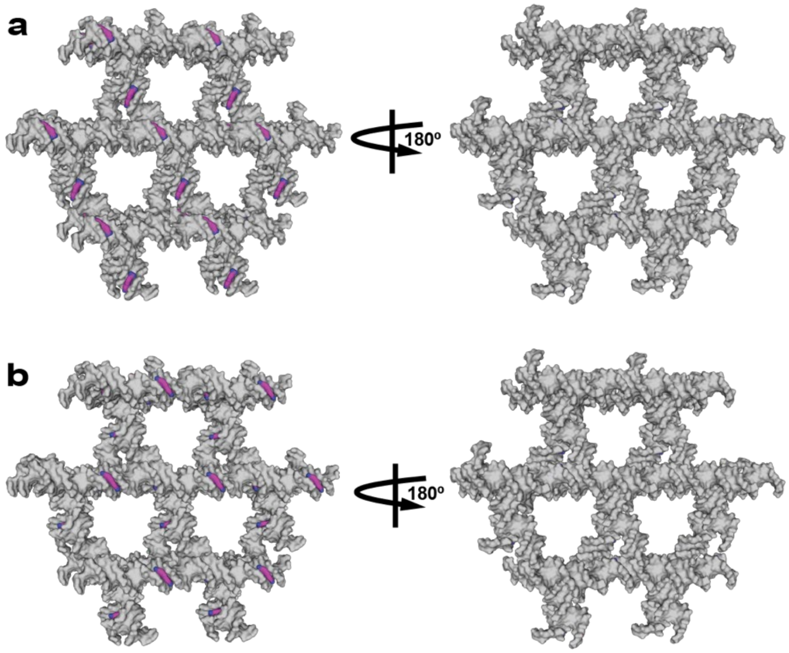

**Supplementary Figure 23. 4x6 crystal lattice packing and orientation of DAPI molecules.** (a) DAPI (purple) Pos1; (b) DAPI (purple) Pos1. Each $P3_2$ lattice is rendered as a surface cross-section of four consecutive layers in the crystal with the AATT minor groove binding sequences at Pos1 (a) and Pos2 (b) in each helical array where each MGB molecule is attached in the structure. The $P3_2$ lattices present all bound MGBs on the front face of the crystal along each constituent duplex in the helical array. When the crystal is oriented 180° no minor grooves on the reverse side contain the bound molecule. The Pos1 and Pos2 lattice results in a 50% occupied crystal. Note: we were unable to determine the structure of the DAPI BP because the crystals did not contain adequate electron density to reliably build the model. The surface of the DAPI (purple carbons) molecules have been surface rendered, with oxygen (red) and nitrogen (blue) also represented.

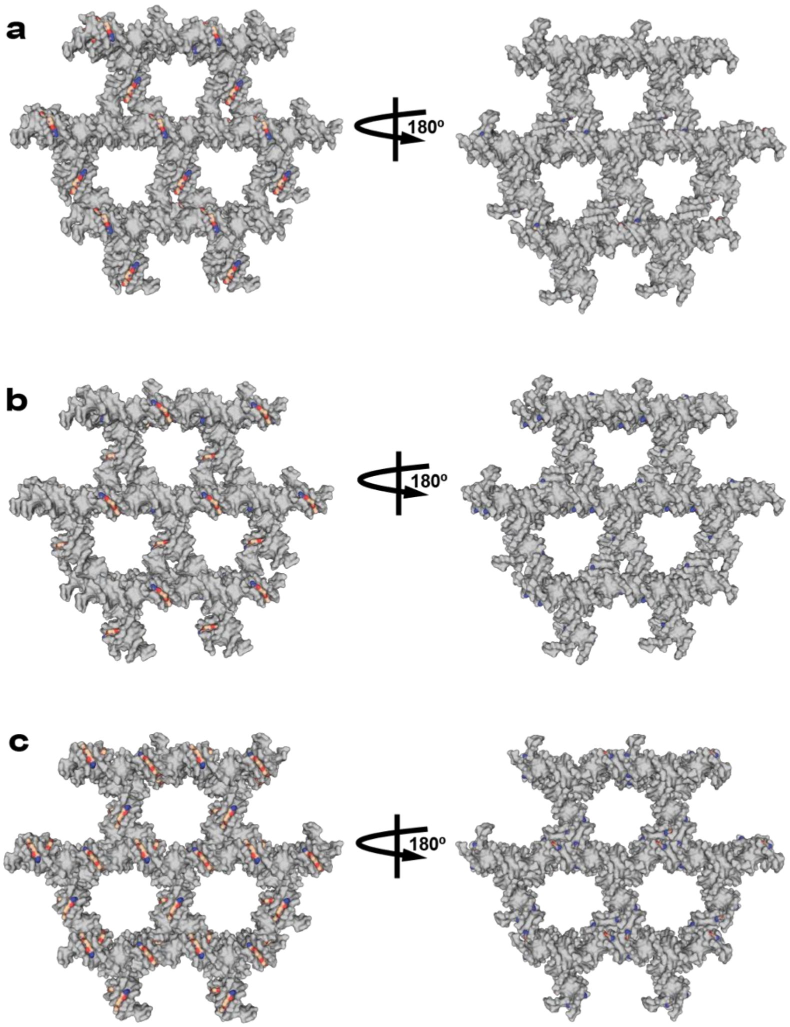

**Supplementary Figure 24. 4x6 crystal lattice packing and orientation of netropsin molecules.** (a) netropsin (tan) Pos1; (b) netropsin (tan) Pos2; (c) netropsin (tan) BP. Each $P3_2$ lattice is rendered as a surface cross-section of four consecutive layers in the crystal with the AATT minor groove binding sequences at Pos1 (a), Pos2 (b), and BP (c) in each helical array where each MGB molecule is attached in the structure. The $P3_2$ lattices present all bound MGBs on the front face of the crystal along each constituent duplex in the helical array. When the crystal is oriented 180° no minor grooves on the reverse side contain the bound molecule. The Pos1 and Pos2 lattice results in a 50% occupied crystal, and the BP crystals yielded 100% occupancy crystals. The surface of the netropsin (tan carbons) molecules have been surface rendered, with oxygen (red) and nitrogen (blue) also represented.

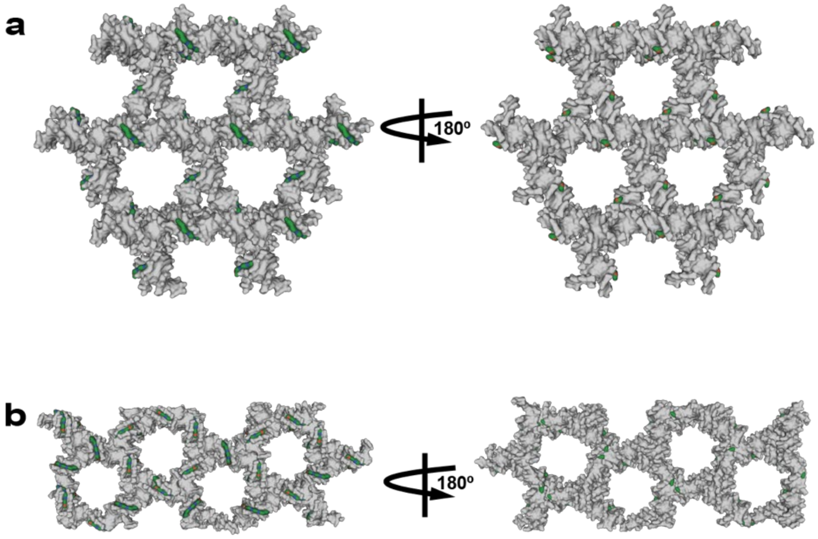

**Supplementary Figure 25. 4x6 crystal lattice packing and orientation of Hoechst molecules.** (a) Hoechst (green) Pos2 with $P3_2$ symmetry; (b) Hoechst (green) BP with *R*3 symmetry. Each lattice is rendered as a surface cross-section of four consecutive layers in the crystal with the AATT minor groove binding sequences at the respective positions in each helical array where each MGB molecule is attached. The $P3_2$ lattices present all bound MGBs on the front face of the crystal along each constituent duplex in the helical array. When the crystal is oriented 180° no minor grooves on the reverse side contain the bound molecule. The Pos2 lattice results in a 50% occupied crystal and the BP crystals contain 100% occupancy. Note: we were unable to determine the structure of the Hoechst Pos1 structure because the crystals did not contain adequate electron density to reliably build the model. It should be noted that the BP Hoechst structure crystallized with both $P3_2$ and *R*3 symmetry in the same buffer; however, only the *R*3 crystals produced adequate electron density to solve the structure. The surface of the Hoechst (green carbons) molecules have been surface rendered, with oxygen (red) and nitrogen (blue) also represented.

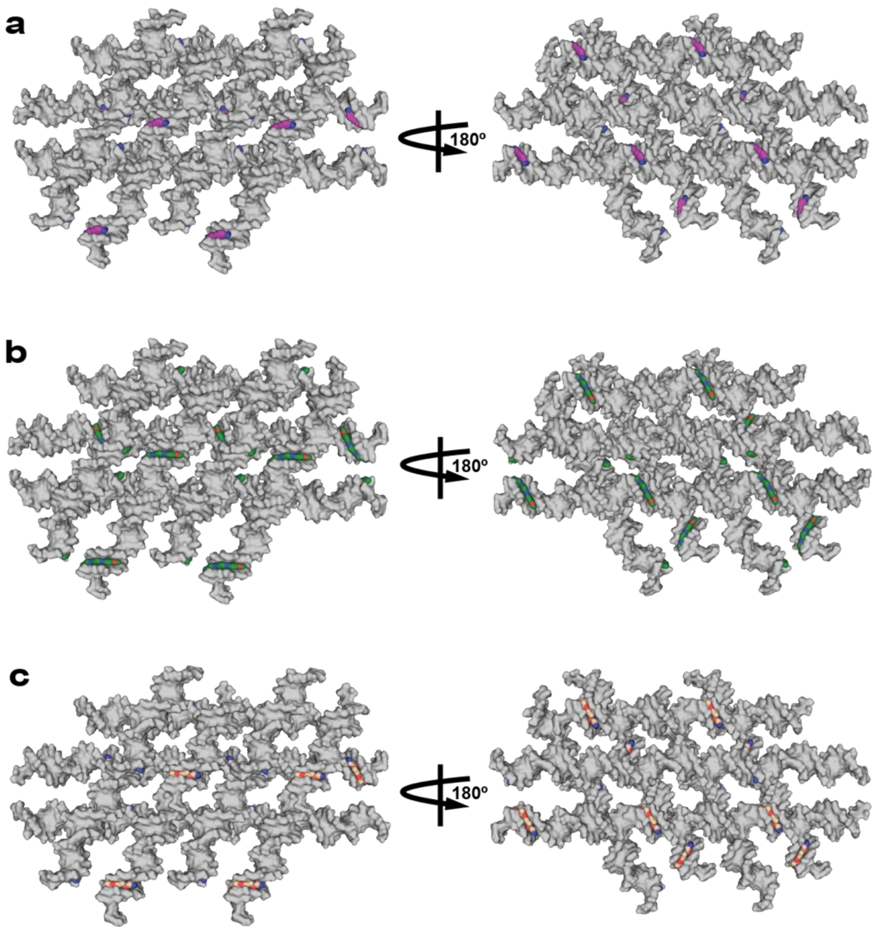

**Supplementary Figure 26. 4x5 Pos1 crystal lattice packing and orientation.** (a) DAPI (purple) Pos1; (b) Hoechst (green) Pos1; (c) netropsin Pos1. Each $P3_221$ lattice is rendered as a surface cross-section between two adjacent layers tethered by Holliday junction crossovers every two full helical turns. The Pos1 minor groove contains the AATT binding sequence to which the target molecules are attached. The $P3_221$ symmetry presents the molecules on the front side of alternating continuous duplexes in the array with the others bound at the Pos1 minor grooves on the opposite side (180°) of their adjacent counterparts. The surface of the DAPI (purple carbons), Hoechst (purple carbons), and netropsin (tan carbons) molecules have been surface rendered, with oxygen (red) and nitrogen (blue) also represented.
